# The NCA-1 and NCA-2 ion channels function downstream of Gq and Rho to regulate locomotion in *C. elegans*

**DOI:** 10.1101/090514

**Authors:** Irini Topalidou, Pin-An Chen, Kirsten Cooper, Shigeki Watanabe, Erik M. Jorgensen, Michael Ailion

## Abstract

The heterotrimeric G protein G_q_ positively regulates neuronal activity and synaptic transmission. Previously, the Rho guanine nucleotide exchange factor Trio was identified as a direct effector of G_q_ that acts in parallel to the canonical G_q_ effector phospholipase C. Here we examine how Trio and Rho act to stimulate neuronal activity downstream of G_q_ in the nematode *Caenorhabditis elegans*. Through two forward genetic screens, we identify the cation channels NCA-1 and NCA-2, orthologs of mammalian NALCN, as downstream targets of the G_q_/Rho pathway. By performing genetic epistasis analysis using dominant activating mutations and recessive loss-of-function mutations in the members of this pathway, we show that NCA-1 and NCA-2 act downstream of G_q_ in a linear pathway. Through cell-specific rescue experiments, we show that function of these channels in head acetylcholine neurons is sufficient for normal locomotion in *C. elegans*. Our results suggest that NCA-1 and NCA-2 are physiologically relevant targets of neuronal G_q_-Rho signaling in *C. elegans*.

## Introduction

Heterotrimeric G protein pathways play central roles in altering neuronal activity and synaptic transmission in response to experience or changes in the environment. G_q_ is one of the four types of heterotrimeric G proteins alpha subunits in animals (Wilkie *et al*. 1992) and is widely expressed in the mammalian brain (Wilkie *et al*. 1991) where it typically acts to stimulate neuronal activity and synaptic transmission (Krause *et al*. 2002; Gamper *et al*. 2004; Coulon *et al*. 2010). These roles are conserved in the nematode *C. elegans*. Unlike mammals which have four members of the G_q_ family, *C. elegans* has only a single G_q_α (Brundage *et al*. 1996). In *C. elegans*, loss-of-function and gain-of-function mutants in the single G_q_α gene *egl-30* are viable but have strong neuronal phenotypes, affecting locomotion, egg-laying, and sensory behaviors (Brundage *et al*. 1996; Lackner *et al*. 1999; Bastiani *et al*. 2003; Matsuki *et al*. 2006; Esposito *et al*. 2010; Adachi *et al*. 2010). We aim to identify the signal transduction pathways and downstream targets by which G_q_ signaling alters neuronal activity.

In the canonical G_q_ pathway, G_q_ activates phospholipase Cβ (PLC) to cleave the lipid phosphatidylinositol 4,5,-bisphosphate (PIP2) into diacylglycerol (DAG) and inositol trisphosphate (IP3), each of which can act as a second messenger. This pathway operates in both worms and mammals, but in both systems, a number of PLC-independent effects of G_q_ have been described, indicating the existence of additional G_q_ signal transduction pathways (Lackner *et al*. 1999; Miller *et al*. 1999; Vogt *et al*. 2003; Bastiani *et al*. 2003; Sánchez-Fernández *et al*. 2014). Using a genetic screen for suppressors of activated G_q_, we identified the Rho guanine nucleotide exchange factor (GEF) Trio as a direct effector of G_q_ in a second major G_q_ signal transduction pathway independent of the canonical PLC pathway (Williams *et al*. 2007). Biochemical and structural studies demonstrated that G_q_ directly binds and activates RhoGEF proteins in both worms and mammals, indicating that this new G_q_ pathway is conserved (Lutz *et al*. 2005, 2007; Williams *et al*. 2007).

G_q_ activation of the Trio RhoGEF leads to activation of the small GTP-binding protein Rho, a major cellular switch that affects a number of cellular processes, ranging from regulation of the cytoskeleton to transcription (Etienne-Manneville and Hall 2002; Jaffe and Hall 2005). In *C. elegans* neurons, Rho has been shown to regulate synaptic transmission downstream of the G_12_-class G protein GPA-12 via at least two pathways, one dependent on the diacylglycerol kinase DGK-1 and one independent of DGK-1 (McMullan *et al*. 2006; Hiley *et al*. 2006). Here we investigate what targets operate downstream of Rho in the G_q_ signaling pathway to regulate neuronal activity. Through two forward genetic screens, we identify the cation channels NCA-1 and NCA-2 (NALCN in mammals) as downstream targets of the G_q_-Rho pathway. The NALCN channel is a relative of voltage-gated cation channels that has been suggested to be a sodium leak channel required for the propagation of neuronal excitation and the fidelity of synaptic transmission (Lu *et al*. 2007; Jospin *et al*. 2007; Yeh *et al*. 2008). However, there is controversy over the current carried by NALCN and whether it is indeed a sodium leak channel (Senatore *et al*. 2013; Senatore and Spafford 2013; Boone *et al*. 2014). It is also unclear how NALCN is gated and what pathways activate the channel. Two studies have shown that NALCN-dependent currents can be activated by G protein-coupled receptors, albeit independently of G proteins (Lu *et al*. 2009; Swayne *et al*. 2009), and another study showed that the NALCN leak current can be activated by low extracellular calcium via a G protein-dependent pathway (Lu *et al*. 2010). Our data presented here suggest that the worm NALCN orthologs NCA-1 and NCA-2 are activated by Rho acting downstream of G_q_ in a linear pathway.

## Materials and Methods

### Strains

Worm strains were cultured and maintained using standard methods (Brenner, 1974). A complete list of strains and mutations used is provided in the strain list (Table S1).

### Isolation of suppressors of activated G_q_

We performed an ENU mutagenesis to screen for suppressors of the hyperactive locomotion of an activated G_q_ mutant, *egl-30(tg26)* (Ailion *et al*. 2014). From approximately 47,000 mutagenized haploid genomes, we isolated 10 mutants that had a fainter phenotype when outcrossed away from the *egl-30(tg26)* mutation. By mapping and complementation testing, we assigned these ten mutants to three genes: three *unc-79* mutants (*yak37*, *yak61*, and *yak73*), six *unc-80* mutants (*ox329*, *ox330*, *yak8*, *yak35*, *yak36*, and *yak56*) and one *nlf-1* mutant (*ox327*). Complementation tests of *ox329* and *ox330* were performed by crossing heterozygous mutant males to *unc-79(e1068)* and *unc-13(n2813) unc-80(ox301)* hermaphrodites. Complementation tests of *yak* alleles were performed by crossing heterozygous mutant males (*m/+*) to *unc-79(e1068)* and *unc-80(ox330)* mutant hermaphrodites. For all crosses, we assessed the fainting phenotype of at least five male cross-progeny by touching animals on the head and scoring whether animals fainted within five seconds. For the crosses to *unc-13 unc-80*, we also scored the fainting phenotype of at least ten Non-Unc-13 hermaphrodites similarly. Control crosses with wild-type males demonstrated that *unc-79/+* and *unc-80/+* heterozygous males and hermaphrodites are phenotypically wild-type and do not show fainting behavior, demonstrating that these mutants are fully recessive.

### Isolation of suppressors of activated G_o_

We first isolated suppressors of the activated G_o_ mutant *unc-109(n499)* by building double mutants of *unc-109(n499)* with the activated G_q_ allele *egl-30(tg26)*. Unlike *unc-109(n499)* homozygotes which are lethal, *egl-30(tg26) unc-109(n499)* homozygotes are viable, but paralyzed and sterile, indicating that activated G_q_ partially suppresses activated G_o_, consistent with the model that these two G proteins act antagonistically. We built a balanced heterozygote strain *egl-30(tg26) unc-109(n499)/egl-30(tg26) unc-13(e51) gld-1(q126)* which has an “unmotivated” phenotype in which the worms move infrequently and slowly. We mutagenized these animals with ENU and screened for F1 progeny that moved better. From a screen of approximately 16,000 mutagenized haploid genomes, we isolated two apparent *unc-109* intragenic mutants, *ox303* and *ox304*. *ox303* is a strong *unc-109* loss-of-function allele, as evidenced by the fact that *egl-30(tg26) unc-109(n499 ox303)/egl-30(tg26) unc-13(e51) gld-1(q126)* mutants resembled *egl-30(tg26)* mutants (i.e. hyperactive). Additionally, *unc-109(n499 ox303)* mutants themselves are hyperactive in an otherwise wild-type background. *ox304*, however, appears to be a partial loss-of-function mutant, because the *egl-30(tg26) unc-109(n499 ox304)/egl-30(tg26) unc-13(e51) gld-1(q126)* mutant moves better than the *egl-30(tg26) unc-109(n499)/egl-30(tg26) unc-13(e51) gld-1(q126)* parent strain, but is not hyperactive like the *egl-30(tg26)* strain. Also, *unc-109(n499 ox304)* homozygote animals on their own are viable but are almost paralyzed and show very little spontaneous movement, with a typical straight posture. However, when stimulated by transfer to a new plate, the *unc-109(n499 ox304)* mutant is surprisingly capable of coordinated movements. This strain was used for mapping and sequencing experiments that demonstrated that *unc-109* is allelic to *goa-1*, encoding the worm G_o_ ortholog (see below). The *unc-109(n499 ox304)* strain was also used as the starting point for a second screen to isolate extragenic suppressors of activated G_o_.

Previously, a screen for suppressors of activated *goa-1* was performed using heat-shock induced expression of an activated *goa-1* transgene (Hajdu-Cronin *et al*. 1999). This screen isolated many alleles of *dgk-1*, encoding diacyclglycerol kinase, but only a single allele of *eat-16*, encoding a regulator of G protein signaling (RGS) protein that negatively regulates G_q_ (Hajdu-Cronin *et al*. 1999), along with mutants in three genes needed for expression of the heat-shock *goa-1* transgene (Hajdu-Cronin *et al*. 2004). Because of the strong bias of this screen for isolating alleles of *dgk-1*, we used the *goa-1(n499 ox304*) strain to perform a screen for suppressors of activated G_o_ that did not involve overexpression of *goa-1* or rely on expression of a heat-shocked induced *goa-1* transgene. We performed ENU mutagenesis of *goa-1(n499 ox304*) and isolated F2 animals that were not paralyzed. From a screen of approximately 24,000 mutagenized haploid genomes, we isolated 17 suppressors, 9 with a relatively stronger suppression phenotype and 8 that were weaker. Of the 9 stronger suppressors, we isolated two alleles of *eat-16* (*ox359*, *ox360*), three alleles of the BK type potassium channel *slo-1* (*ox357*, *ox358*, *ox368*), one allele of the gap junction innexin *unc-9* (*ox353*), one gain-of-function allele in the ion channel gene *nca-1* (*ox352*), and two mutants that were mapped to chromosomal regions distinct from the other mutations listed above, but not further characterized (*ox356*, *ox364*). The G_o_ suppressors we isolated are consistent with the established model that activation of GOA-1 activates the RGS EAT-16, which inhibits G_q_ EGL-30 (Hajdu-Cronin *et al*. 1999). Reduced G_q_ activity leads to reduced activation of the NCA cation channels, which can be reversed by an activating mutation in NCA-1. Previously, it has been shown that lack of the depolarizing NCA cation currents can be suppressed by a compensatory loss of the hyperpolarizing potassium current from SLO-1 (Kasap *et al*. 2017), or by loss of the gap junction proteins UNC-9 or UNC-7 (Sedensky and Meneely 1987; Morgan and Sedensky 1995; Bouhours *et al*. 2011). This is consistent with our isolation of *slo-1* and *unc-9* mutants as suppressors of activated G_o_, since activation of G_o_ also leads to reduced NCA activity.

### Mapping and cloning *nlf-1(ox327)*

We mapped the *ox327* mutation using single nucleotide polymorphisms (SNPs) in the Hawaiian strain CB4856 as described (Davis *et al*., 2005). The *ox327* mutation was mapped to an approximately 459 kb region on the left arm of the X chromosome between SNPs on cosmids F39H12 and C52B11 (SNPs F39H12[4] and pkP6101). This region included 74 predicted protein-coding genes. We injected cosmids spanning this region and found that injection of cosmid F55A4 rescued the *ox327* mutant phenotype. We performed RNAi to the genes on this cosmid in the *eri-1(mg366) lin-15(n744)* strain that has enhanced RNAi and found that RNAi of the gene F55A4.2 caused a weak fainter phenotype. We sequenced F55A4.2 in the *ox327* mutant and found a T to A transversion mutation in exon 1, leading to a premature stop codon at amino acid C59. We also rescued the *ox327* mutant with a transgene carrying only F55A4.2, confirming the gene identification. We subsequently obtained a deletion allele *tm3631* that has fainter and G_q_ suppression phenotypes indistinguishable from *ox327*. F55A4.2 was given the gene name *nlf-1* (Xie *et al*. 2013).

We obtained six independent *nlf-1* cDNAs that were predicted to be full-length: yk1105g4, yk1159a2, yk1188d11, yk1279a1, yk1521f8, and yk1709b10. Restriction digests suggested that all six were of the same size. We sequenced yk1159a2 and yk1279a1 and both gave the same *nlf-1* exon-intron structure, which differed from the gene-structure on Wormbase WS253 in several ways: *nlf-1* is 4 bp shorter at the 3’ end of exon 5, and has a new 154 bp exon (now exon 6) not predicted on Wormbase. *nlf-1* consists of 8 exons and is predicted to encode a protein of 438 amino acids (Figure 3A). This is identical to the gene structure reported independently (Xie *et al*. 2013). Both sequenced cDNAs had 5’UTRs of 64 bp, with yk1159a2 (but not yk1279a1) being trans-spliced to the SL1 splice leader, and 3’UTRs of 424 bp (yk1279a1) or 429 bp (yk1159a2). The yk1279a1 cDNA was mutation-free and was cloned into a Gateway entry vector for use in rescue experiments. The full-length sequence of the yk1279a1 *nlf-1* cDNA was deposited in GenBank under accession # KX808524.

### Mapping and cloning *unc-109(n499)*

*unc-109* was shown to be allelic to *goa-1*. First, we performed SNP mapping of both *unc-109(n499)* and its intragenic revertant *unc-109(n499 ox303)*, using the Hawaiian strain CB4856 as described (Davis *et al*. 2005). These experiments mapped *unc-109* to an approximately 1 Mb region in the middle of chromosome I between SNPs on cosmids D2092 and T24B1 (SNPs CE1-15 and T24B1[1]). A good candidate in this region was *goa-1*. We confirmed that *unc-109* was indeed *goa-1* by sequencing the three *unc-109* mutants: the gain-of-function allele *n499* carries a point mutation that leads to an R179C missense mutation, affecting a conserved arginine residue shown to be important for the GTPase activity of G proteins (Coleman *et al*. 1994); the partial loss-of-function allele *ox304* carries a point mutation leading to a W259R missense mutation; and the strong loss-of-function allele *ox303* carries a one basepair deletion that leads to a premature stop 32 amino acids from the C-terminal.

### Molecular biology and transgenes

A complete list of constructs is provided in the plasmid list (Table S2). Most of the constructs were made using the three slot multisite Gateway system (Invitrogen). For C3 transferase constructs, a promoter, an FRT-mCherry-FRT-GFP cassette (pWD178), and the *C. botulinum* C3 transferase-*unc-54* 3’UTR (cloned into a Gateway entry vector from plasmid QT#99) were combined into the pDEST R4-R3 destination vector. For *nlf-1* tissue-specific rescue constructs, a promoter, the *nlf-1* coding sequence (genomic DNA or cDNA), and a C-terminal GFP tag were cloned along with the *unc-54* 3’UTR into the pDEST R4-R3 destination vector. Promoters used were *nlf-1p* (5.7 kb upstream of the ATG), *rab-3p* (all neurons), *unc-17p* (acetylcholine neurons), *unc-17Hp* (head acetylcholine neurons) (Hammarlund *et al*. 2007), *acr-2p* (acetylcholine motor neurons), *unc-17*β*p* (acetylcholine motor neurons) (Charlie *et al*. 2006), and *glr-1p* (glutamate-receptor interneurons). Extrachromosomal arrays were made by standard transformation methods (Mello *et al*. 1991). Constructs of interest were injected at 10 ng/μl with marker and carrier DNAs added to make a final total concentration of at least 100 ng/μl. For most constructs, we isolated multiple independent insertions that behaved similarly. C3 transferase extrachromosomal arrays were integrated into the genome using X-ray irradiation (4000 rads). Integrated transgenes were mapped to chromosomes and outcrossed twice before further analysis.

### Locomotion assays

We performed two different assays to measure locomotion. Body bend assays measured the rate of locomotion. Radial locomotion assays measured the radial distance animals moved from a point in a given unit of time, which provides a combined measurement of different aspects of the locomotion phenotype including the rate of locomotion, waveform, and frequency of reversals. Both types of assays were performed on 10 cm plates seeded with thin lawns of OP50 bacteria. These plates were prepared by seeding 1.5 ml of stationary phase OP50 bacteria to evenly cover the entire plate surface, and then growing the bacteria for two days at room temperature. Plates were stored at 4° for up to one month before being used. For body bend assays, first-day adult worms were picked to an assay plate, allowed to rest for 30 seconds, and then body bends were counted for one minute. A body bend was defined as the movement of the worm from maximum to minimum amplitude of the sine wave (Miller *et al*. 1999). To minimize variation, all animals in a body bend experiment were assayed on the same plate. For radial locomotion assays, five to eight first-day adults were picked together to the center of a plate to begin the assay (time 0). Positions of the worms were marked on the lid of the plate every ten minutes for up to forty minutes. Following the assay, the distance of each point to the center was measured. For most strains, radial distances did not increase after the first ten minutes of the assay and all data presented here are for the ten-minute time point. Analysis of the data at later time points leads to the same conclusions. For all locomotion assays, the experimenter was blind to the genotypes of the strains being assayed.

For Rho inhibition experiments (Figure 1), expression of C3 transferase (C3T) was induced by FLP-mediated recombination. Expression of FLP was induced by heat shock for 1 hr at 34°C, plates were returned to room temperature, and animals were scored for locomotion 4 hrs after the end of the heat shock period. For heat-shock induction of activated Rho (Figure 8), worms were heat shocked for 1 hr at 34°C, returned to room temperature, and scored for locomotion 2 hrs after the end of the heat-shock period.

**Figure 1.**
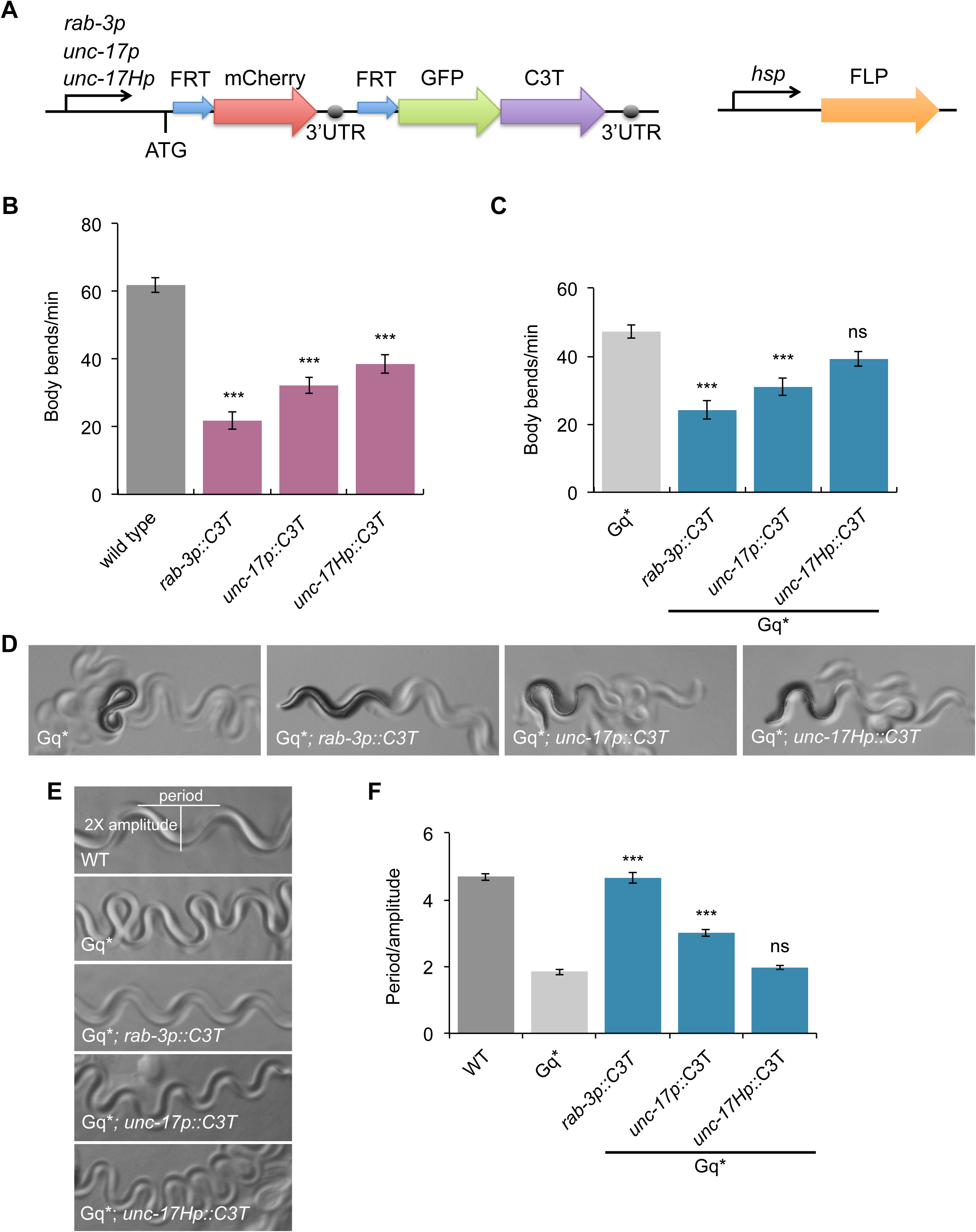
Inhibition of Rho in neurons suppresses an activated G_q_ mutant. (A) Schematic of the FLP/FRT system we used for temporal and spatial expression of the Rho inhibitor C3 transferase (C3T). The transgene in the ‘‘off’’ configuration expresses the mCherry reporter under the control of the promoter sequence but terminates transcription in the *let-858* 3’UTR upstream of GFP-C3T. Expression of the FLP recombinase is induced by heat shock and leads to recombination between the FRT sites and deletion of the intervening *mCherry::let-858* 3’UTR fragment. This leads to transcription of GFP-C3T in the cells driven by the adjacent promoter. We used the pan-neuronal promoter *rab-3p,* acetylcholine neuron promoter *unc-17p,* and head acetylcholine neuron promoter *unc-17Hp.* (B) Inhibition of Rho in adult neurons causes a locomotion defect. Inhibition of Rho by C3 transferase (C3T) in all adult neurons (*rab-3p, oxIs412* transgene), acetylcholine neurons (*unc-17p, oxIs414* transgene) or head acetylcholine neurons (*unc-17Hp, oxIs434* transgene) reduces the locomotion rate of wild type animals. ***, P<0.001, Dunnett’s test. Error bars = SEM; n = 19-20. (C) Inhibition of Rho in all adult neurons (*rab-3p, oxIs412* transgene) or acetylcholine neurons (*unc-17p*, *oxIs414* transgene) led to a significant reduction in the rate of locomotion of the activated G_q_ mutant *egl-30(tg26)* (here written Gq*). Inhibition of Rho in head acetylcholine neurons (*unc-17Hp*, *oxIs434* transgene) led to a small decrease in Gq* locomotion that was not statistically significant. ***, P<0.001; ns, not significant, P>0.05, Dunnett’s test. Error bars = SEM; n = 10-20. (D) Photos of first-day adult worms. The coiled body posture of the activated G_q_ mutant *egl-30(tg26)* (Gq*) is suppressed by Rho inhibition in all neurons (*rab-3p::C3T*). (E) Photos of worm tracks. (F) Quantification of the track waveform. The high-amplitude waveform of the activated G_q_ mutant *egl-30(tg26)* (Gq*) is suppressed by Rho inhibition in all adult neurons (*rab-3p::C3T*) and partially suppressed by Rho inhibition in acetylcholine neurons (*unc-17p::C3T*). Rho inhibition in head acetylcholine (*unc-17Hp::C3T*) neurons did not suppress the waveform of Gq*. ***, P<0.001; ns, not significant, P>0.05, Bonferroni test. All comparisons are to Gq*. Error bars = SEM; n = 5.

### Fainting assays

Backward fainting times were measured by touching a worm on the head with a worm pick to stimulate movement, and measuring the time to faint with a stopwatch. Forward fainting time was measured following a touch on the tail. Fainting was defined by an abrupt stop of movement along with a characteristic straightening of the head (Figure 6A). Alternatively, we touched worms on the head or tail with a pick and counted the number of body bends until the worm fainted. If a worm moved 10 body bends without fainting, we stopped the assay.

### Waveform quantification

To quantify the track waveform, first-day adult animals were placed on an OP50 plate and allowed to move forward for a few seconds. We then imaged each animal’s tracks using a Nikon SMZ18 microscope with the DS-L3 camera control system. Track pictures were taken at 40X and were processed using ImageJ. Period and 2X amplitude were measured using the line tool. For each worm, five period/amplitude ratios were averaged. Five individual worms were used per experiment.

The exaggerated waveform of *egl-30(tg26)* mutants is also characterized by the head of the worm occasionally crossing over the body, leading the worm to form a figure-eight shape (Figure 2A). This phenotype was quantified by placing animals on OP50 plates and allowing them to move forward, counting the number of times the head crossed over the body in one minute.

**Figure 2.**
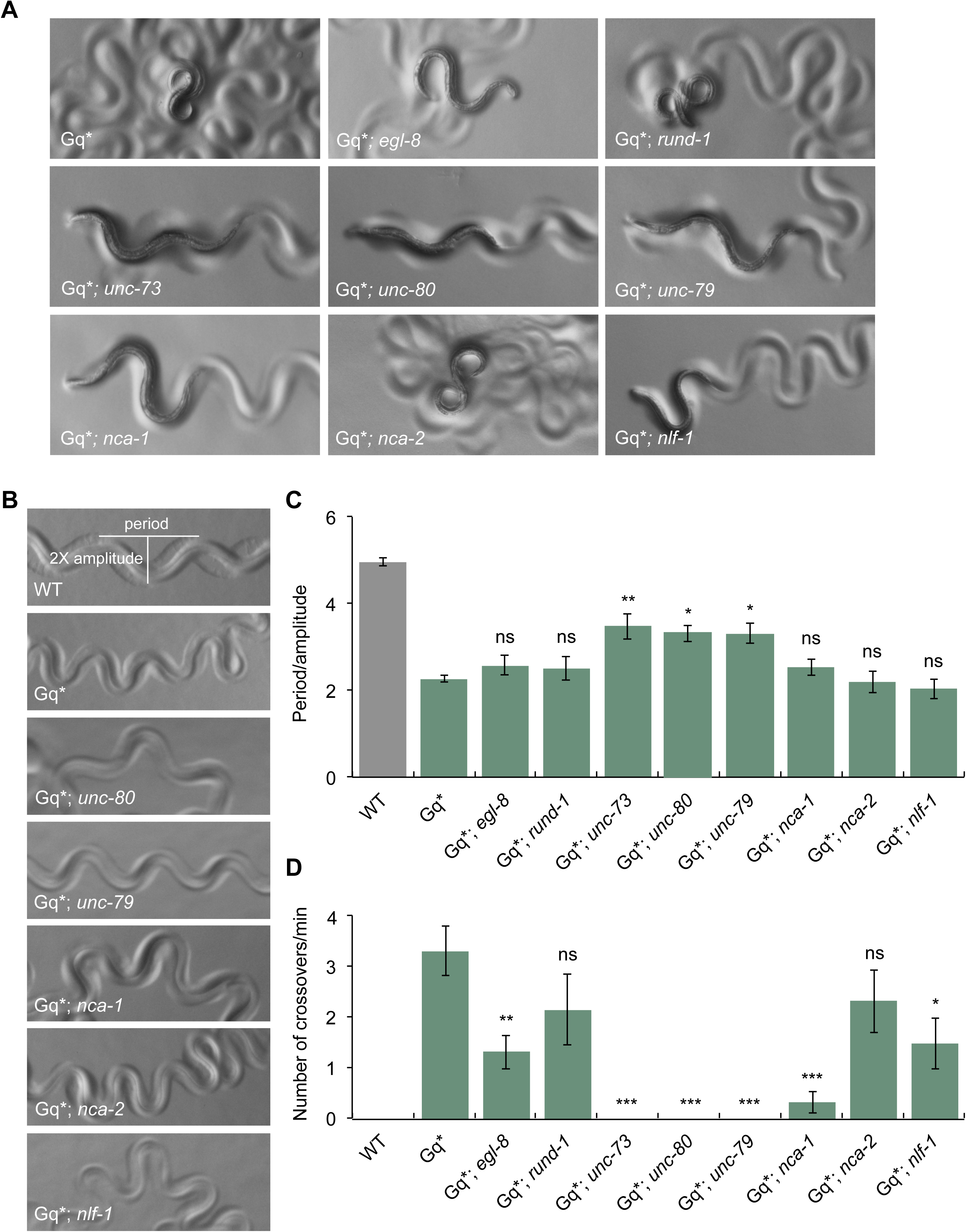
Mutations affecting the NCA-1 and NCA-2 channels suppress the high-amplitude waveform of the activated G_q_ mutant *egl-30(tg26)* (Gq*). The *unc-73(ox317)*, *unc-79(yak37)* and *unc-80(ox330)* mutations suppress the high-amplitude waveform of Gq*. The *nca-1(gk9)* and *nlf-1(ox327)* mutations partially suppress the high-amplitude waveform of Gq* as measured by the crossover assay. Mutations in the canonical G_q_ pathway such as *egl-8(ox333)* partially suppress and genes required for dense-core vesicle biogenesis such as *rund-1(tm3622)* do not significantly suppress the high-amplitude waveform of Gq*. The *nca-2(gk5)* mutation does not suppress Gq*. (A) Photos of first-day adults. (B) Photos of worm tracks. The photo of wild type tracks is the same as the one shown in Figure 3C. (C) Quantification of the track waveform. The wild-type waveform data are the same data shown in Figures 3D, 6C, and 9D. (D) Quantification of the worm’s head crossing its body (“crossovers”). ***, P<0.001; **, P<0.01; *, P<0.05; ns, not significant, P>0.05, Bonferroni test. All comparisons are to Gq*. Error bars = SEM; n = 5 (panel C); n = 6 (panel D).

**Figure 3.**
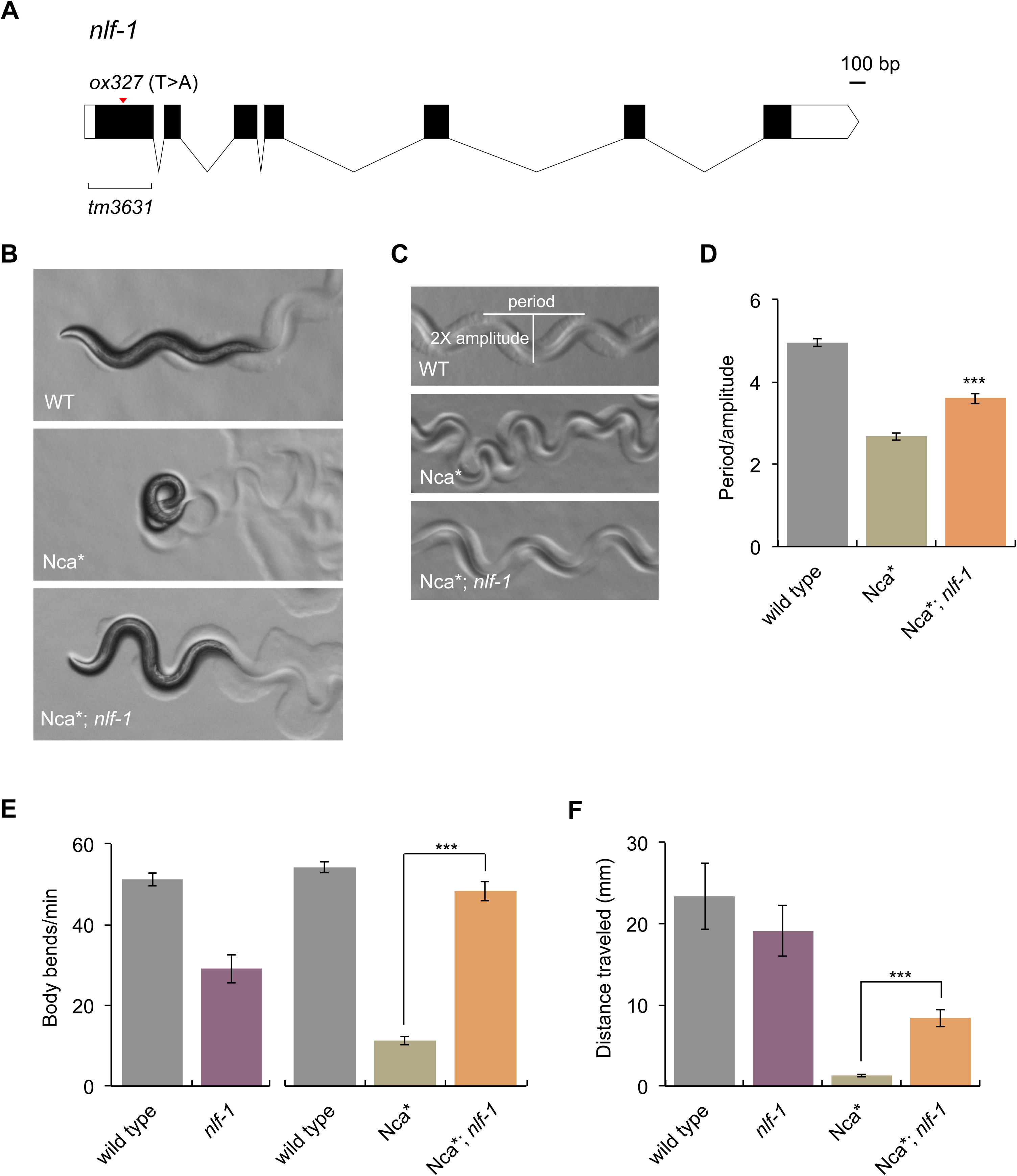
The *nlf-1(ox327)* mutation suppresses the coiled posture, high-amplitude waverform, and locomotion defect of the activated NCA-1 mutant *nca-1(ox352*) (Nca*). (A) Gene structure of *nlf-1*. Black boxes show coding segments. White boxes show 5’ and 3’ untranslated regions (UTRs). The *ox327* mutation leads to a premature stop codon at C59. The *tm3631* deletion removes most of the 5’UTR and first exon. (B) Photos of first-day adult worms. (C) Photos of worm tracks. The photo of wild type tracks is the same as the one shown in Figure 2B. (D) Quantification of track waveform. The high-amplitude waveform of Nca* is suppressed by *nlf-1.* The wild-type data are the same data shown in Figures 2C, 6C, and 9D. (E) Body bend assay. The wild-type data shown in the left graph are the same as shown in Figure 7A, right graph. (F) Radial locomotion assay. Statistics in D: ***, P<0.001, two-tailed unpaired t test. Comparisons are to Nca*. Error bars = SEM, n = 5. Statistics in E and F: ***, P<0.001, two-tailed Mann-Whitney test. Error bars = SEM, n = 4-10 (panel E), n = 8-26 (panel F).

### Imaging and image analysis

Worms were mounted on 2% agarose pads and anesthetized with sodium azide. Images were obtained using a Zeiss Pascal confocal microscope. For quantitative imaging of NCA-1::GFP and NCA-2::GFP (Figure S1), Z-stack projections of the nerve ring axons on one side of the animal were collected and quantified in ImageJ as described (Jospin *et al*. 2007). Dissecting microscope photographs of first-day adult worms were taken at 50X using a Nikon SMZ18 microscope equipped with a DS-L3 camera control system.

### Statistics

P values were determined using GraphPad Prism 5.0d (GraphPad Software). Normally distributed data sets with multiple comparisons were analyzed by a one-way ANOVA followed by a Bonferroni or Tukey posthoc test to examine selected comparisons or by Dunnett’s test if all comparisons were to the wild type control. Non-normally distributed data sets with multiple comparisons were analyzed by a Kruskal-Wallis nonparametric ANOVA followed by Dunn’s test to examine selected comparisons. Pairwise data comparisons were analyzed by a two-tailed unpaired t test for normally distributed data or by a two-tailed Mann-Whitney test for non-normally distributed data.

### Reagent and data availability

Strains and plasmids are shown in Table S1 and Table S2 and are available from the *Caenorhabditis* Genetics Center (CGC) or upon request. The full-length sequence of the yk1279a1 *nlf-1* cDNA was deposited in GenBank under accession # KX808524. The authors state that all data necessary for confirming the conclusions presented in the article are represented fully within the article and Supplemental Material.

## Results

### Inhibition of Rho suppresses activated G_q_

Two pieces of data suggested that G_q_ may regulate locomotion in *C. elegans* through activation of the small G protein Rho. First, both the hyperactive locomotion and tightly wound body posture of the activated G_q_ mutant *egl-30(tg26)* are suppressed by loss-of-function mutations in the *unc-73* RhoGEF Trio, an activator of Rho (Williams *et al*. 2007). Second, expression of activated Rho as a transgene causes worms to adopt a posture characterized by a tightly coiled, high-amplitude waveform (McMullan *et al*. 2006), reminiscent of the waveform of activated G_q_ worms (Bastiani *et al*. 2003; Ailion *et al*. 2014). To determine directly whether G_q_ signals through Rho, we tested whether Rho inhibition suppresses an activated G_q_ mutant.

In *C. elegans,* there is a single gene encoding Rho (*rho-1*). Loss of *rho-1* causes numerous pleiotropic developmental phenotypes and embryonic lethality (Jantsch-Plunger *et al*. 2000), making it difficult to study Rho function using classical loss-of-function approaches. As an alternative method of inactivating Rho, we employed the *Clostridium botulinum* C3 transferase (C3T) which ADP-ribosylates Rho and has been widely used as a Rho inhibitor (Aktories *et al*. 2004). C3T has strong substrate specificity for Rho and has only weak activity towards other Rho family members like Rac and Cdc42 (Just *et al*. 1992). Though effects of C3T on targets other than Rho cannot be absolutely excluded, a study in *C. elegans* found that C3T, *rho-1* RNAi, and expression of a dominant-negative form of Rho all caused similar defects in P cell migration (Spencer *et al*. 2001). To bypass the developmental roles and study Rho function in the adult nervous system, we expressed the C3T only in adult neurons by using the FLP recombinase/FRT system (Davis *et al*. 2008). In this system, temporal control of C3T is achieved through induction of FLP via heat-shock. FLP in turn promotes recombination between FRT sites to lead to expression of C3 transferase in specific neurons (Figure 1A). We expressed C3T in the following classes of neurons: all neurons (*rab-3p*), acetylcholine neurons *(unc-17p*), head acetylcholine neurons (*unc-17Hp*), and acetylcholine motor neurons (*unc-17*β*p*). mCherry fluorescence confirmed expression in the expected neurons and GFP expression was used to monitor induction of C3T following FLP-mediated recombination.

Inhibition of *rho-1* in adult neurons caused a decreased locomotion rate (Figure 1B). This effect was greatest when Rho was inhibited in all neurons, but Rho inhibition in acetylcholine subclasses of neurons also led to slower locomotion. Thus, Rho acts in multiple classes of neurons to promote locomotion in adult worms. In the absence of heat-shock, all strains showed normal wild-type rates of locomotion and did not express any detectable GFP, indicating that these transgenes do not provide leaky expression of C3T in the absence of heat-shock. This confirms that Rho acts post-developmentally in mature neurons to regulate locomotion behavior (McMullan *et al*. 2006).

To determine whether Rho acts downstream of G_q_ signaling, we crossed the C3 transgenes into the background of the activated G_q_ mutant *egl-30(tg26)*. Inhibition of *rho-1* in all adult neurons strongly suppressed the high-amplitude waveform, body posture, and hyperactive locomotion of the activated G_q_ mutant (Figure 1, C-F). Inhibition of *rho-1* in acetylcholine neurons suppressed the hyperactivity of activated G_q_ (Figure 1C), but only weakly suppressed the high-amplitude waveform (Figure 1, D-F). Thus, *rho-1* exhibits genetic interactions consistent with a role in the G_q_ signaling pathway in both acetylcholine neurons and additional neurons.

### Mutations in NCA channel subunits suppress activated G_q_

What acts downstream of Rho in the G_q_ signal transduction pathway? We screened for suppressors of activated G_q_ and found mutants in three categories: (1) the canonical G_q_ pathway (such as the PLC *egl-8*); (2) the RhoGEF *unc-73* (Williams *et al*. 2007), and (3) genes that affect dense-core vesicle function (*e.g. unc-31*, *rab-2*, *rund-1*) (Ailion *et al*. 2014; Topalidou *et al*. 2016). Mutations in *unc-73* strongly suppress the high-amplitude waveform of the activated G_q_ mutant (Figure 2, A and C), but mutations in the canonical G_q_ pathway or in genes that affect dense-core vesicle function only weakly suppress the high-amplitude waveform of the activated G_q_ mutant (Figure 2). Thus, strong suppression of the high-amplitude waveform of activated G_q_ may be a specific characteristic of mutations in the Rho pathway, and we hypothesized that downstream targets of Rho in this pathway would also suppress the high-amplitude waveform.

To identify possible downstream targets of Rho in the G_q_ pathway, we examined other mutants isolated from our screen for suppression of the high-amplitude waveform of activated G_q_ (Figure 2). When crossed away from the activating G_q_ mutation, several of these mutants had a “fainter” phenotype. Fainter mutants respond to a touch stimulus by moving away, but abruptly stop, that is “faint”, after only a few body bends. The fainter phenotype has been observed only in mutants that reduce the function of the NCA-1 and NCA-2 ion channels (Humphrey *et al*. 2007; Jospin *et al*. 2007; Yeh *et al*. 2008). We found that all our G_q_ suppressors that strongly suppressed the high-amplitude waveform and had a fainter phenotype were mutants in either *unc-79* or *unc-80*, two genes required for function of the NCA channels (Humphrey *et al*. 2007; Jospin *et al*. 2007; Yeh *et al*. 2008). We also isolated a single mutant in the gene *nlf-1* that also gave fainters after outcrossing away from the activated G_q_ mutation, but did not strongly suppress the high-amplitude waveform of the activated G_q_ mutant (Figure 2). *unc-79* and *unc-80* mutants have a strong fainter phenotype equivalent to that of a double mutant in *nca-1* and *nca-2*, two genes that encode pore-forming subunits of the NCA channels in *C. elegans* (Humphrey *et al*. 2007; Jospin *et al*. 2007; Yeh *et al*. 2008). Like *unc-79* or *unc-80*, an *nca-1 nca-2* double mutant suppressed the activated G_q_ mutant. Additionally, *nca-1* on its own partially suppressed activated G_q_, but *nca-2* did not (Figure 2). This suggests that although *nca-1* and *nca-2* are only redundantly required for normal worm locomotion, channels containing the NCA-1 pore-forming subunit have a larger role in transducing G_q_ signals than NCA-2 channels.

### Cloning and characterization of *nlf-1(ox327)*

In addition to the previously known NCA channel subunits *unc-79* and *unc-80*, we also isolated the *ox327* mutant in a gene that had not been previously characterized at the time of our study. We cloned *ox327* by single nucleotide polymorphism (SNP) mapping, RNAi and transgenic rescue experiments (see Materials and Methods), showing that it carries an early stop mutation in the gene *nlf-1*. We sequenced two *nlf-1* cDNAs and found that its exon-intron structure differed from the gene structure predicted on Wormbase (Figure 3A, see Materials and Methods for details). *nlf-1* was independently cloned by others (Xie *et al*. 2013).

*nlf-1* encodes an endoplasmic reticulum-localized protein probably involved in proper assembly of the NCA channel, since *nlf-1* mutants had reduced expression levels of GFP-tagged NCA-1 and NCA-2 (Xie *et al*. 2013). We also found that *nlf-1* is required for normal axonal levels of both GFP-tagged NCA-1 and NCA-2 in the nerve ring (Figure S1). Additionally, an *nlf-1* mutation suppressed both the coiled posture and slow locomotion of an activated *nca-1* mutant (Figure 3, B-F), demonstrating that *nlf-1* is important for NCA-1 function.

*nlf-1* mutants have a weaker fainter phenotype than mutants of *unc-79* or *unc-80*, or the *nca-1 nca-2* double mutant (Figure 4, A and B). The *nlf-1* fainting phenotype differs in two ways from those of the stronger fainting mutants. First, *nlf-1* mutants take a longer time to faint following stimulation (Figure 4, A and B). Second, while the strong fainting mutants show a similarly strong fainting phenotype in either the forward or backward direction, *nlf-1* mutants faint reliably in the backward direction but take much longer and have more variable fainting in the forward direction (Figure 4, A and B), suggesting that *nlf-1* mutations cause a partial loss of function of the NCA channels. To determine how *nlf-1* interacts with NCA mutants, we built *nlf-1(ox327)* double mutants with *unc-79, unc-80*, *nca-1*, and *nca-2*. The *nlf-1* mutation did not enhance the *unc-79* or *unc-80* fainter phenotype, suggesting that these mutants act in the same pathway to control fainting (Figure 4, C and D). However, *nca-1* strongly enhanced the *nlf-1* fainter phenotype, but *nca-2* did not significantly enhance *nlf-1* (Figure 4, A and B). Neither *nca-1* nor *nca-2* single mutants have a fainter phenotype on their own (Humphrey *et al*. 2007), but the fact that an *nca-1 nlf-1* double mutant has a strong fainter phenotype suggests that either *nca-1* contributes more than *nca-2* for normal locomotion or that *nlf-1* specifically perturbs function of *nca-2*. Our data presented below are more consistent with the possibility that *nca-1* contributes more than *nca-2* to wild-type locomotion behavior.

**Figure 4.**
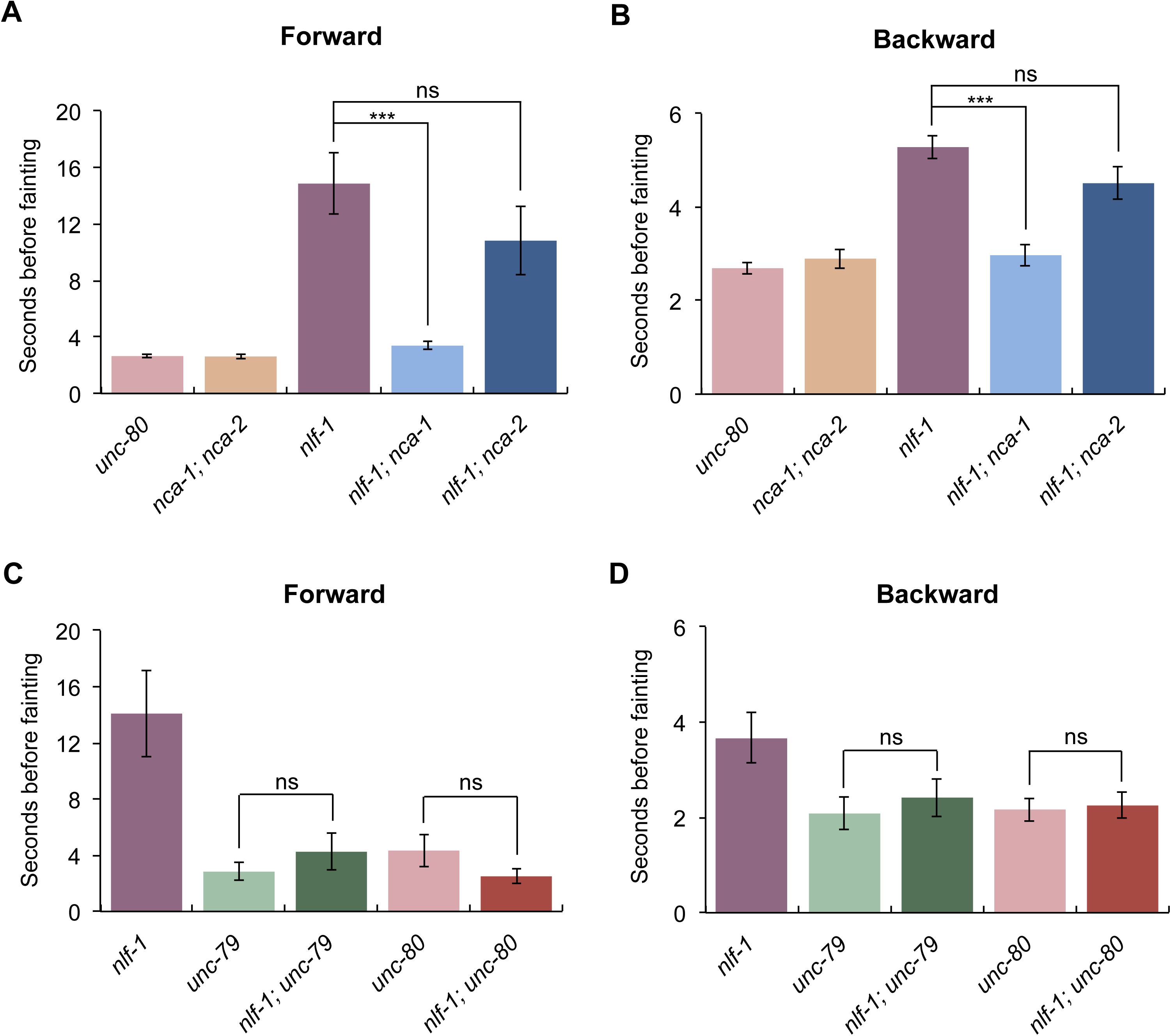
*nlf-1* mutants are weak fainters. The *nlf-1(ox327)* mutant is a weaker fainter in the forward direction (A) than the backward direction (B), but *nlf-1(ox327)* has a weaker fainter phenotype than either *unc-80(ox330)* or the *nca-2(gk5)*; *nca-1(gk9)* double mutant. Additionally, *nlf-1* mutants are enhanced by a mutation in *nca-1*, but not *nca-2*. Wild-type, *nca-1(gk9)*, and *nca-2(gk5)* mutant animals are not shown in the figure because they do not faint (0/10 animals fainted in a one minute period). Also, *nlf-1(ox327)* mutants do not enhance the strong forward (C) or backward (D) fainting phenotypes of *unc-79(e1068)* and *unc-80(ox330)* mutants. ***, P<0.001; ns, not significant, P>0.05, Dunn’s test. Error bars = SEM; n = 10-20.

We determined the cellular expression pattern of *nlf-1* by fusing its promoter to GFP. *nlf-1p::GFP* was expressed in most or all neurons, but was not detected in other tissues (Figure 5A). This agrees with the expression pattern reported elsewhere (Xie *et al*. 2013). To determine the neuronal focus of the fainter phenotype, we performed rescue experiments in which we determined whether an *nlf-1* mutant could be rescued by expression of a wild-type *nlf-1* cDNA under the control of neuron-specific promoters. Expression of *nlf-1(+)* in all neurons (using the *rab-3* promoter) or acetylcholine neurons (using the *unc-17* promoter) fully rescued the *nlf-1* mutant fainter phenotype (Figure 5B). Expression in acetylcholine motor neurons (using the *acr-2* or *unc-17*β promoters) did not rescue the fainter phenotype, but expression driven by a head-specific derivative of the *unc-17* promoter (*unc-17Hp*) fully rescued the fainter phenotype, indicating that the action of *nlf-1* in head acetylcholine neurons is sufficient to prevent fainting (Figure 5B).

Previously, it was reported that expression of *nlf-1* in premotor interneurons is sufficient to rescue the *nlf-1* mutant fainter phenotype (Xie *et al*. 2013). However, we found that expression of *nlf-1* in premotor interneurons using the *glr-1* promoter (*glr-1p*) had only a weak effect on the *nlf-1* mutant fainter phenotype and did not fully restore wild-type locomotion (Figure 5C). Fainting behavior was difficult to score in the *glr-1* promoter-rescued *nlf-1* mutant animals because they had sluggish movement and stopped frequently, though generally not with the suddenness and characteristic posture typical of fainters. Though we could blindly score the rescue of fainting in these animals by eye, we saw only weak rescue of the *nlf-1* mutant by *glr-1* promoter expression in our quantitative fainting assays and the effect was not statistically significant (Figure 5C). When we instead measured fainting as the percentage of animals that fainted within ten body bends, we did see a marginally significant rescue by *glr-1* promoter expression (backward fainting: *nlf-1 =* 92%, *nlf-1; glr-1p::nlf-1(+)* = 68%, P=0.0738, Fisher’s exact test; forward fainting: *nlf-1* = 80%, *nlf-1; glr-1p::nlf-1(+)* = 48%, P=0.0378, Fisher’s exact test). We may be underestimating the rescue of the fainting phenotype by *glr-1* promoter expression due to the difficulty distinguishing the frequent pausing from true fainting. Nevertheless, rescue of the *nlf-1* mutant is clearly stronger by expression in head acetylcholine neurons using the *unc-17H* promoter (Figure 5B). Though our data seem to contradict the previous study reporting that *nlf-1* acts in premotor interneurons (Xie *et al*. 2013), there are several possible explanations. First, like our data, the data in the previous study in fact showed only partial rescue of fainting behavior by expression in premotor interneurons (Xie *et al*. 2013). Second, the premotor interneuron promoter combination used in the previous study (*nmr-1p* + *sra-11p*) leads to expression in several other head interneurons that may contribute to the phenotype. Third, it is possible that rescue is sensitive to expression level and that different levels of expression were achieved in the two studies, leading to different levels of rescue. We conclude that NLF-1 acts in head acetylcholine neurons, including the premotor interneurons, to promote sustained locomotion in the worm. Consistent with this, the premotor command interneurons have recently been shown to use acetylcholine as a neurotransmitter (Pereira *et al*. 2015).

**Figure 5.**
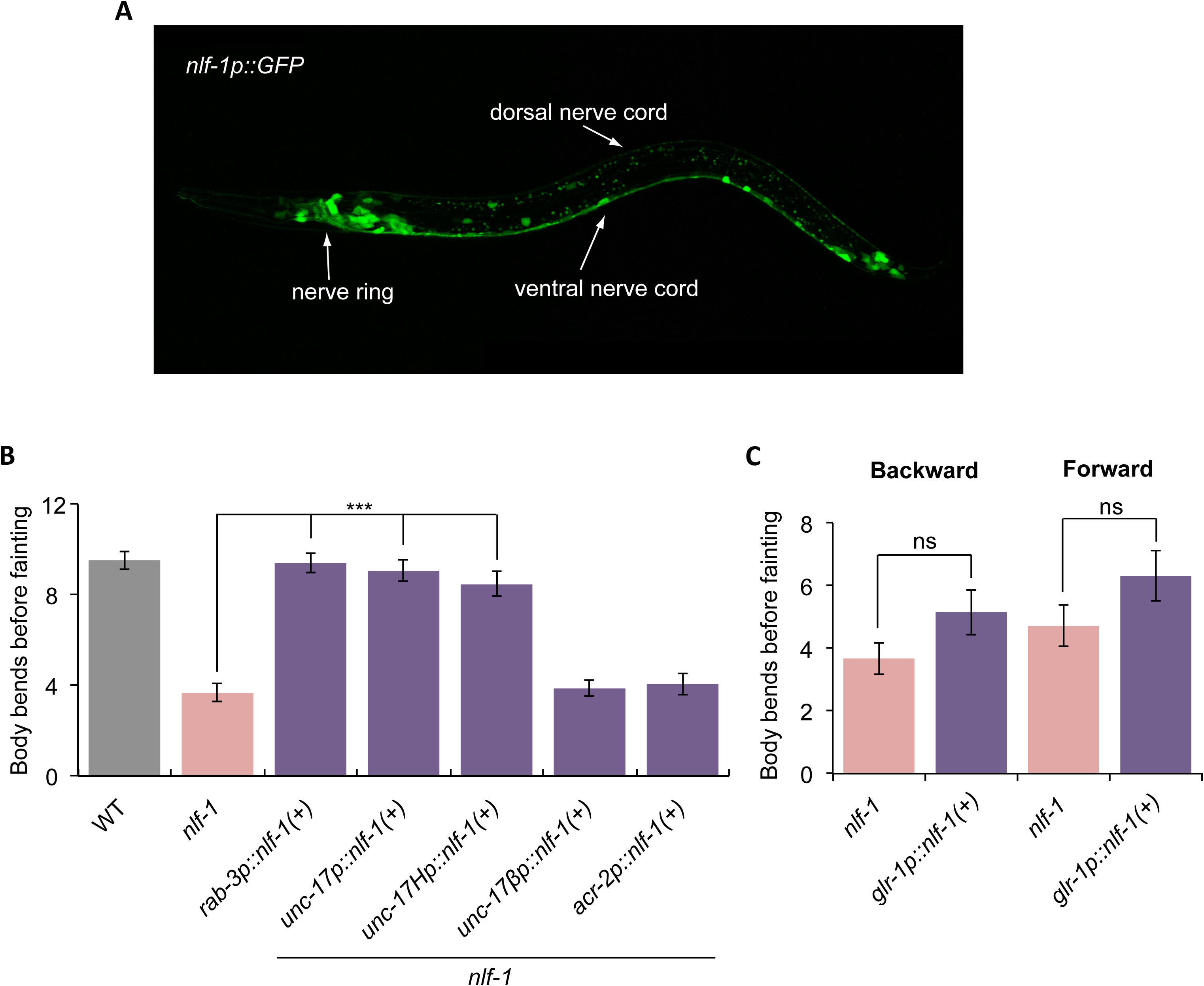
*nlf-1* acts in head acetylcholine neurons to control locomotion. (A) *nlf-1* is expressed widely in the nervous system. A fusion of the *nlf-1* promoter to GFP (transgene *oxEx1144*) is expressed throughout the nervous system, including head and tail neurons, and motor neurons in the ventral nerve cord. No expression is seen in non-neuronal tissues. An L1 stage larval animal is shown with its head to the left. (B,C) The *nlf-1* cDNA was expressed in an *nlf-1(ox327)* mutant background using the following promoters: *rab-3* (all neurons, *oxEx1146* transgene), *unc-17* (acetylcholine neurons, *oxEx1149* transgene), *unc-17H* (a derivative of the *unc-17* promoter that lacks the enhancer for ventral cord expression and thus expresses in head acetylcholine neurons and occasionally a few tail neurons, *oxEx1155* transgene), *unc-17*β (a derivative of the *unc-17* promoter that expresses only in ventral cord acetylcholine neurons, *oxEx1323* transgene), *acr-2* (ventral cord acetylcholine motor neurons, *oxEx1151* transgene), and *glr-1* (interneurons including premotor command interneurons, *oxEx1152* transgene). (B) Expression riven by the *rab-3*, *unc-17*, and *unc-17H* promoters rescued the fainting phenotype of an *nlf-1* mutant in the backward direction. ***, P<0.001, Dunn’s test. Error bars = SEM; n = 15. (C) Expression driven by the *glr-1* promoter partially rescued the fainting phenotype of an *nlf-1* mutant in either the backward or forward direction, though the effect was not statistically significant (ns, P>0.05, two-tailed Mann-Whitney tests. Error bars = SEM; n = 25). However, assays of this strain were complicated by slow movement and frequent pauses that were hard to distinguish from true fainting behavior (see text for details).

### Mutations in NCA channel subunits suppress activated Rho

To determine whether NCA mutants act downstream of Rho, we took advantage of an activated Rho mutant (G14V) expressed specifically in the acetylcholine neurons (McMullan *et al*. 2006). Like an activated G_q_ mutant, this activated Rho mutant has an exaggerated waveform. We built double mutants of the activated Rho mutant with mutations in *unc-79*, *unc-80*, *nlf-1* and mutations in the NCA channel genes *nca-1* and *nca-2*. The *nca-1* and *nca-2* genes are redundant for the fainter phenotype because neither mutant has a fainter phenotype individually, but the double mutant has a fainter phenotype indistinguishable from *unc-79* and *unc-80* mutants (Humphrey *et al*. 2007; Jospin *et al*. 2007; Yeh *et al*. 2008). We found that the high-amplitude waveform caused by activated Rho was strongly suppressed in *unc-80* mutants, as well as in *nca-1 nca-2* double mutants (Figure 6). In both cases, the resulting double or triple mutants had a fainter phenotype like the *unc-80* or *nca-1 nca-2* mutants on their own. By contrast, the high-amplitude waveform of activated Rho was incompletely suppressed by a mutation in *nlf-1*, consistent with the *nlf-1* mutation causing only a partial loss of NCA channel function (Figure 6). Reciprocally, activated Rho suppressed the weak fainter phenotype of an *nlf-1* mutant, again because Rho can act on NCA even in the absence of *nlf-1*. Additionally, a mutation in *nca-1* also partially suppressed the high-amplitude waveform of activated Rho, but a mutation in *nca-2* did not suppress the high-amplitude waveform (Figure 6). Thus, channels containing the NCA-1 pore-forming subunit have a larger role than NCA-2 channels in transducing Rho signals, similar to the interaction of NCA-1 and G_q_ signaling.

**Figure 6.**
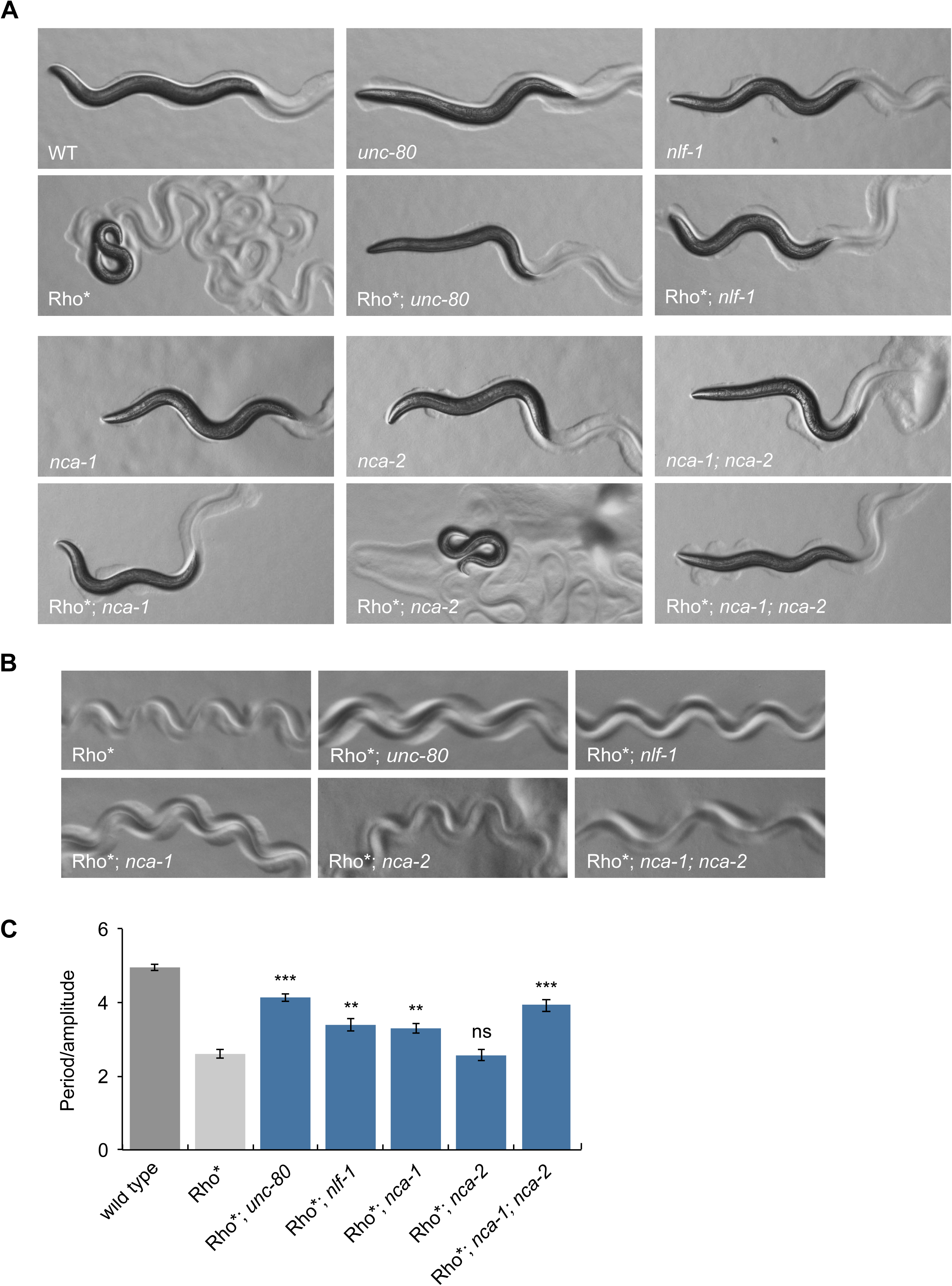
Mutations in NCA channel subunits suppress the high-amplitude waveform of animals expressing activated Rho. Animals expressing an activated Rho mutant (G14V) in acetylcholine neurons (*nzIs29* transgene, written as Rho*) have a high-amplitude waveform. Mutations that eliminate NCA-1 and NCA-2 channel function (*unc-80(ox330)* or *nca-2(gk5)*; *nca-1(gk9)*) strongly suppress the high-amplitude waveform of activated Rho and convert the activated Rho mutants to fainters. Mutations in *nlf-1(ox327)* and *nca-1(gk9)* strongly but incompletely suppress the high-amplitude waveform of activated Rho and these double mutants do not faint. Mutations in *nca-2(gk5)* do not suppress the high-amplitude waveform of activated Rho. Mutants in *unc-80*, *nlf-1*, and the *nca-2 nca-1* double mutant have the typical fainter posture characterized by a straightened anterior part of the body. (A) Photos of first-day adult worms. (B) Photos of worm tracks. (C) Quantification of track waveform. The wild-type data are the same data shown in Figures 2C, 3D, and 9D. ***, P<0.001; **, P<0.01; ns, not significant, P>0.05, Bonferroni test. All comparisons are to Rho*. Error bars = SEM; n = 5.

Because an activated Rho mutant has slow locomotion (Figure 7A) and *unc-79*, *unc-80*, and *nca-1 nca-2* double mutants also have slow locomotion, it is difficult to determine whether these NCA channel mutants suppress the locomotion phenotype of activated Rho in addition to its waveform. We performed radial locomotion assays that provide a combined measurement of several aspects of the locomotion phenotype, including the rate of movement, frequency of reversals, and amplitude of the waveform (see Materials and Methods). By these assays, mutations in *unc-80* or *nca-1 nca-2* lead to only small increases in the radial distance traveled by an activated Rho mutant and an *nca-2* mutant had no effect (Figure 7B). However, a mutation in *nlf-1* much more strongly increased the radial distance traveled by an activated Rho mutant. Because the *nlf-1* mutant is not as slow on its own, we could also directly assay its effect on the rate of locomotion of an activated Rho mutant by counting the number of body bends per minute. An *nlf-1* mutation strongly increased the rate of locomotion of an activated Rho mutant (Figure 7A). In fact, the *nlf-1* double mutant with activated Rho had a faster rate of locomotion than either activated Rho or *nlf-1* on its own, similar to the effect of *nlf-1* on the locomotion of an activated *nca-1* mutant (Figure 3E). Additionally, a mutation in *nca-1* strongly increased the locomotion rate of the activated Rho mutant, but a mutation in *nca-2* had no effect (Figure 7A), further supporting the idea that NCA channels consisting of the NCA-1 subunit act downstream of G_q_ and Rho in this pathway.

**Figure 7.**
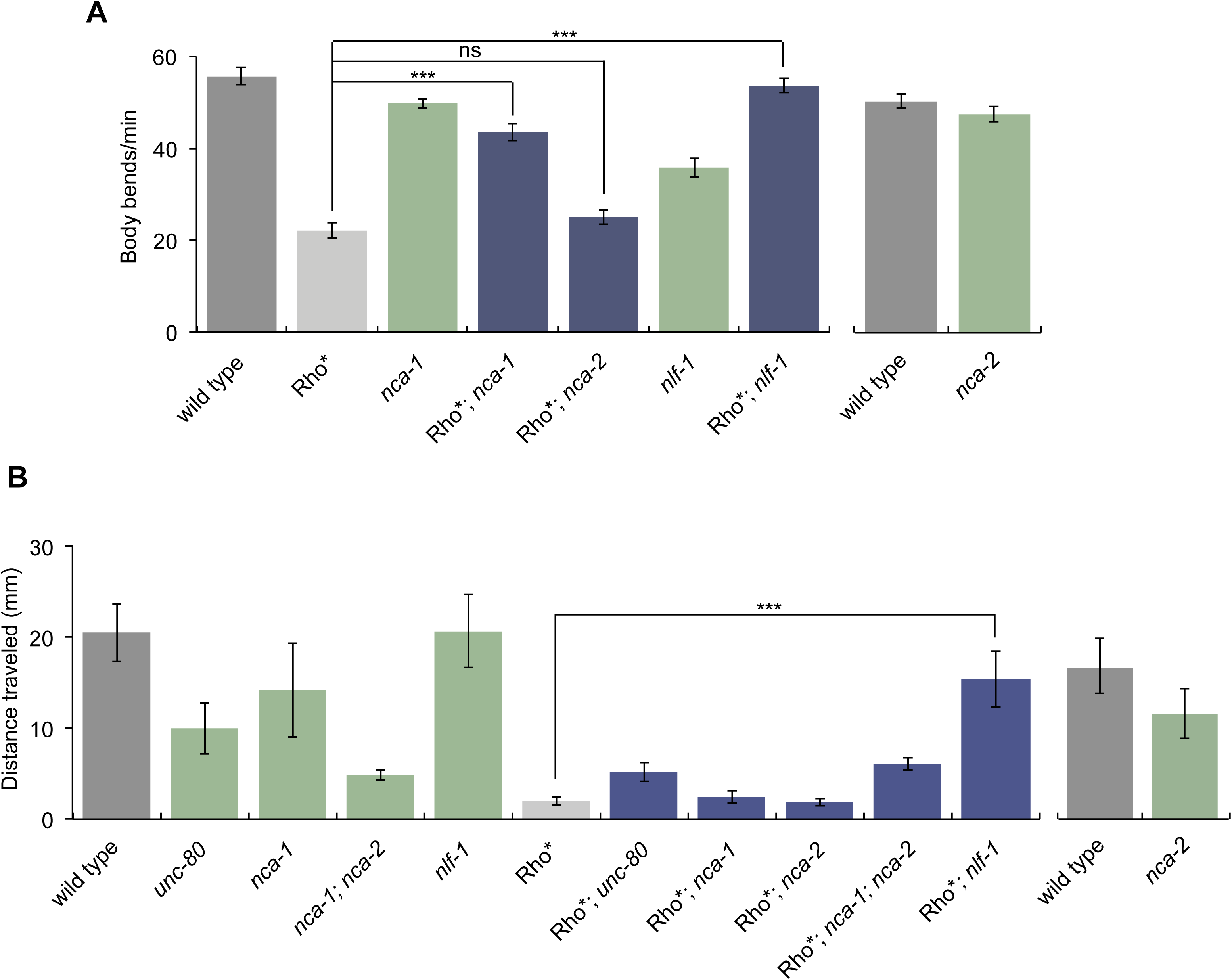
Mutations in *nlf-1* and *nca-1* suppress the locomotion defect of animals expressing activated Rho. (A) Animals expressing an activated Rho mutant (G14V) in acetylcholine neurons (*nzIs29* transgene, written as Rho*) have a slow locomotion rate as measured by the number of body bends. The *nlf-1(ox327)* and *nca-1(gk9)* mutations, but not *nca-2(gk5)*, strongly suppress the slow locomotion rate of activated Rho. The wild-type data shown in the right graph are the same as shown in Figure 3E, left graph. (B) Animals expressing an activated Rho mutant (G14V) in acetylcholine neurons (Rho*) have reduced locomotion as measured in radial locomotion assays. Mutations that eliminate function of the NCA channels (*unc-80(ox330)* or *nca-2(gk5)*; *nca-1(gk9)*) increase the radial distance traveled by activated Rho mutants, but the effect is subtle since *unc-80* and *nca-2; nca-1* mutants have reduced locomotion on their own. The *nlf-1(ox327)* mutation that partially reduces function of the NCA channels strongly suppresses the locomotion defect of activated Rho. In the particular radial locomotion experiment shown here, only five Rho\**; nca-1* animals were assayed and they had a mean radial distance traveled of 2.4 mm, very similar to the mean radial distance of the Rho* single mutant (2.0 mm, n=18). However, in an independent experiment, the Rho\**; nca-1* double mutant had a mean radial distance of 12.2 mm (n=10) as compared to 3.5 mm for the Rho* single mutant (n=10). Thus, it is likely that *nca-1* does indeed suppress *Rho\** in radial locomotion assays, but gave a negative result in the presented assay due to low n, with the likely explanation that these particular Rho*; *nca-1* animals moved outward and returned to the center. ***, P<0.001; ns, not significant, P>0.05, Bonferroni test. Error bars = SEM; n = 10-16 (panel A); n = 5-18 (panel B).

Rho regulates worm locomotion independently of effects on development, as demonstrated by the fact that heat-shock induction of an activated Rho transgene in adults leads to a high-amplitude waveform similar to that seen in worms that express activated Rho in acetylcholine neurons (McMullan *et al*. 2006). Consistent with the idea that NCA-1 acts downstream of Rho, mutations in the fainter genes *unc-80* and *nlf-1* suppress the high-amplitude waveform phenotype of heat-shock induced activated Rho (Figure 8, A-C). Additionally, *nlf-1* also suppresses the locomotion defect of heat-shock induced activated Rho (Figure 8D). Thus, Rho regulates worm locomotion via the NCA channels by acting in a non-developmental pathway that operates in adult neurons.

**Figure 8.**
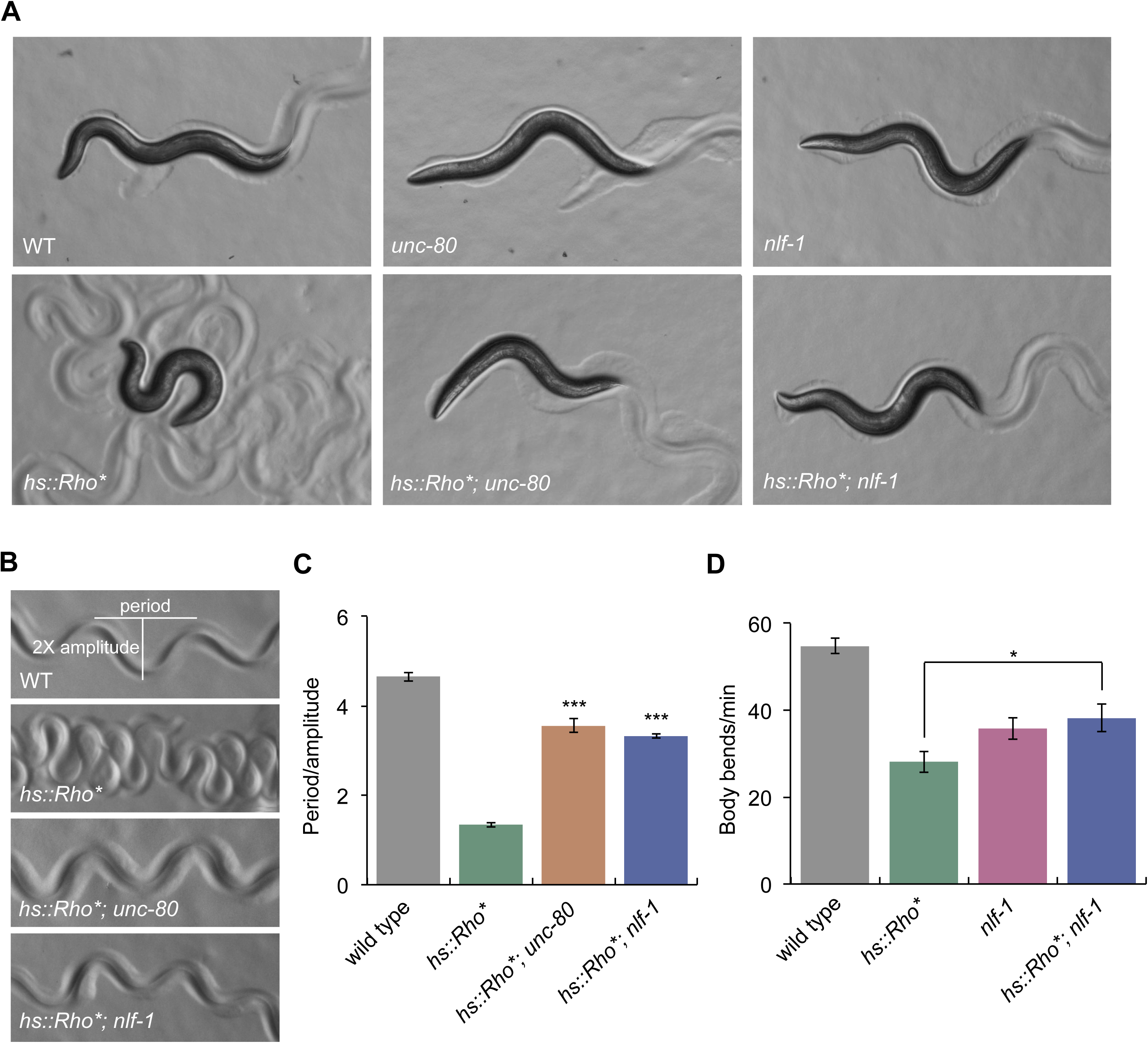
Mutations in NCA channel subunits suppress the high-amplitude waveform and locomotion of adult animals expressing activated Rho. Animals expressing an activated Rho mutant (G14V) only in adults by heat-shock induced expression (*nzIs1* transgene, *hs::Rho\**) have a high-amplitude waveform. The *unc-80(ox330)* and *nlf-1(ox327)* mutations strongly suppress the high-amplitude waveform of activated Rho. *unc-80*, but not *nlf-1*, makes the activated Rho animals faint. (A) Photos of first-day adult worms. (B) Photos of worm tracks. (C) Quantification of track waveform. (D) Animals expressing an activated Rho mutant (G14V) only in adults by heat-shock induced expression (*nzIs1* transgene, *hs::Rho\**) have a reduced locomotion rate. The *nlf-1(ox327)* mutation suppresses the reduced locomotion rate of activated Rho. Statistics in C: ***, P<0.001, Bonferroni test. Error bars = SEM; n = 5. Statistics in D: *, P=0.0157, two-tailed unpaired t test. Error bars = SEM; n = 18-30.

### A dominant NCA-1 mutation suppresses activated G_o_

In *C. elegans*, the G_q_ pathway is opposed by signaling through the inhibitory G_o_ protein GOA-1 (Hajdu-Cronin *et al*. 1999; Miller *et al*. 1999). Thus, loss-of-function mutants in *goa-1* are hyperactive and have a high-amplitude waveform, similar to the gain-of-function G_q_ mutant *egl-30(tg26)*. We found that the uncloned dominant mutant *unc-109(n499)* which is paralyzed and resembles loss-of-function mutants in *egl-30* (Park and Horvitz 1986) carries an activatin 1mutation in *goa-1* (see Materials and Methods). The *goa-1(n499)* mutant is paralyzed as a heterozygote, and is lethal as a homozygote (Park and Horvitz 1986). We performed a screen for suppressors of *goa-1(n499)* and isolated a partial intragenic suppressor, *goa-1(n499 ox304*) (Materials and Methods). *goa-1(n499 ox304)* homozygote animals are viable but are almost paralyzed and show very little spontaneous movement, with a straight posture and low-amplitude waveform (Figure 9, C and D). However, when stimulated, the *goa-1(n499 ox304)* mutant is capable of slow coordinated movements (Figure 9, A and B).

**Figure 9.**
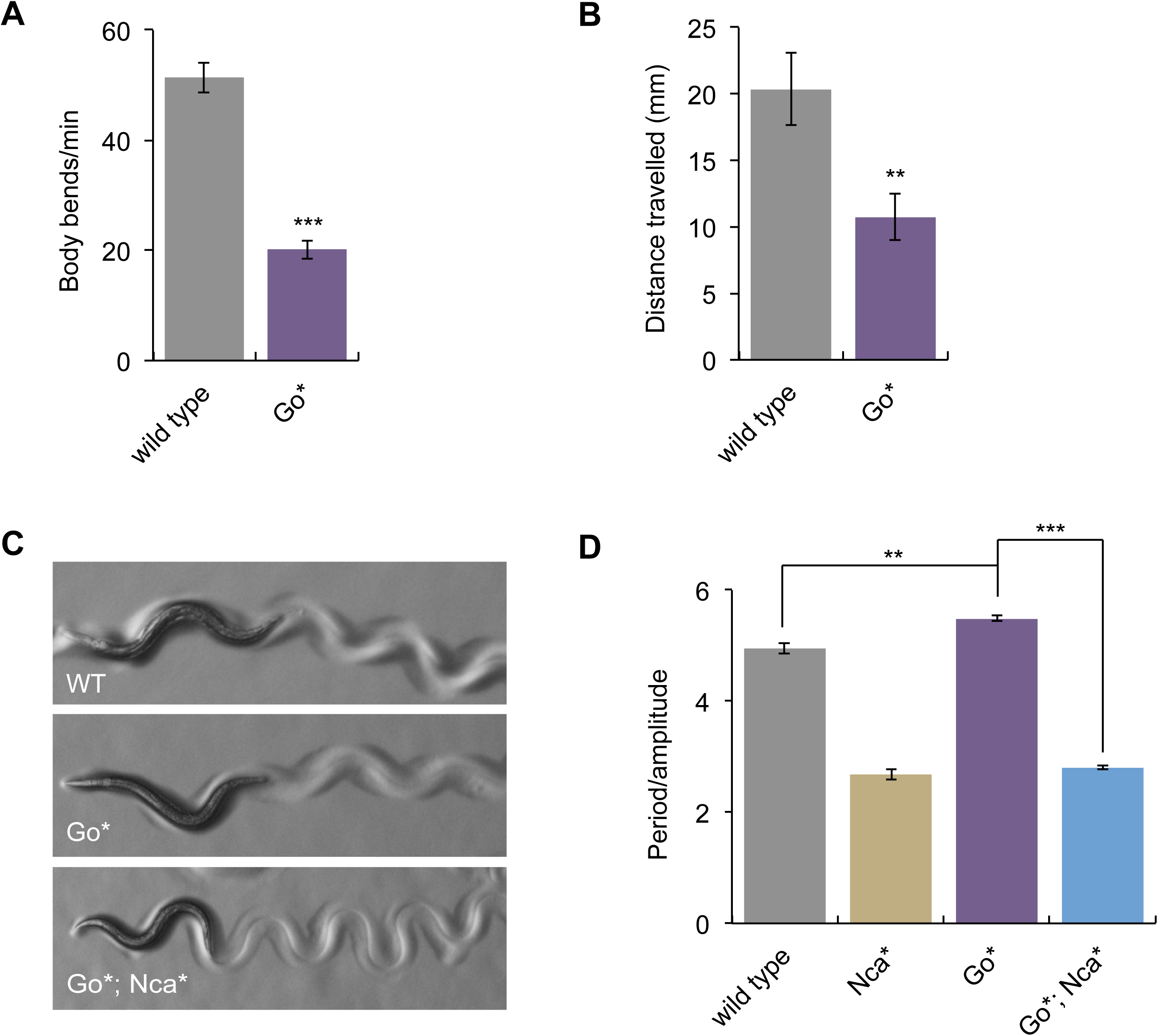
An activated NCA-1 mutant suppresses the straight posture and low-amplitude waveform of an activated GOA-1 mutant. The *goa-1(n499 ox304)* mutant (written as Go*) has a strong movement defect as shown by a body bend assay (A) and a radial locomotion assay (B).***, P<0.001; **, P<0.01, two-tailed unpaired t-tests. Error bars = SEM; n = 10 (panel A); n = 18 (panel B). Also the *goa-1(n499 ox304)* mutant has a relatively straight posture (C) and low-amplitude waveform (D) that is suppressed by the *nca-1(ox352)* mutation (written as Nca*). The wild-type waveform data are the same data shown in Figures 2C, 3D, and 6C. ***, P<0.001; **, P<0.01, Bonferroni test. Comparisons are to Go*. Error bars = SEM; n = 5-6.

We mutagenized the *goa-1(n499 ox304*) strain and screened for animals that were not paralyzed or did not have a straight posture. Among the suppressors, we isolated a mutant (*ox352*) that was found to be a dominant gain-of-function mutation in the *nca-1* gene (Bend *et al*. 2016). The *nca-1(ox352)* mutant has a coiled posture and high-amplitude waveform reminiscent of the activated G_q_ and activated Rho mutants (Figure 3B). Furthermore*, nca-1(ox352*) suppresses the straight waveform of the activated G_o_ mutant *goa-1(n499 ox304)* (Figure 9, C and D). Thus, activation of NCA-1 suppresses activated G_o_ whereas loss of NCA-1 suppresses activated G_q_, both consistent with the model that G_o_ inhibits G_q_ and that NCA-1 is a downstream effector of the G_q_ pathway.

## Discussion

In this study, we identify the NCA-1 and NCA-2 ion channels as downstream effectors of the heterotrimeric G proteins G_o_ and G_q_. Previously, it has been demonstrated that activation of G_o_α GOA-1 activates the RGS protein EAT-16, which in turn inhibits G_q_α (Hajdu-Cronin *et al*. 1999; Miller *et al*. 1999). Loss of *goa-1* leads to hyperactive worms (Mendel *et al*. 1995; Ségalat *et al*. 1995). Here we identified an activated mutation in *goa-1* that causes animals to be paralyzed, closely resembling null mutations in the G_q_α gene *egl-30* (Brundage *et al*. 1996). Suppressors of activated G_o_ include loss-of-function mutations in the RGS EAT-16, and gain-of-function mutations in the NCA-1 channel, suggesting that activated G_o_ inactivates G_q_, and could thereby indirectly inactivate the NCA-1 cation channel.

Further genetic epistasis data indicate that G_q_ activates the RhoGEF Trio and the small GTPase Rho. Rho then acts via an unknown mechanism to activate the NCA-1 and NCA-2 ion channels, which are required for normal neuronal activity and synaptic transmission in *C. elegans* (Jospin *et al*. 2007; Yeh *et al*. 2008; Xie *et al*. 2013; Gao *et al*. 2015). Thus, this work identifies a new genetic pathway from G_q_ to an ion channel that regulates neuronal excitability and synaptic release: G_q_ → RhoGEF → Rho → NCA channels (Figure 10). The NCA channels have not been previously identified as effectors of the G_q_ pathway, so this pathway may give insight into how the NCA channels are activated or regulated.

**Figure 10.**
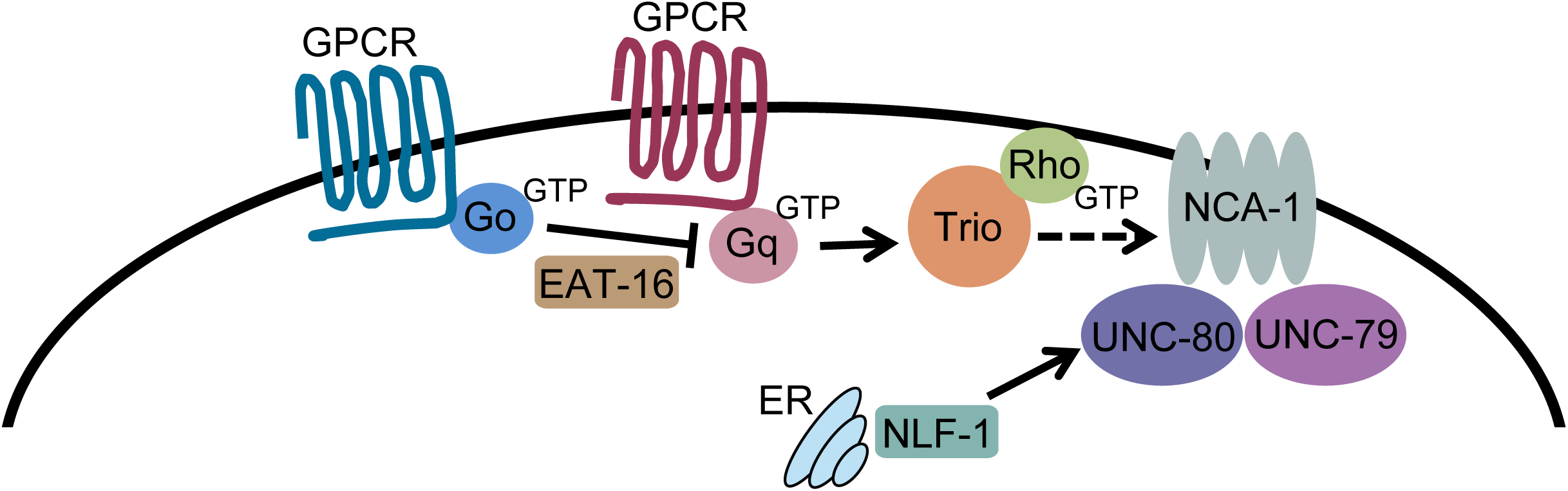
Model. G_q_ activates NCA-1 in a linear pathway via the RhoGEF Trio and small G protein Rho. G_q_ activation of Trio and Rho is direct (solid arrow). Rho activation of the NCA-1 channel is likely to be indirect (dashed arrow). G_o_ inhibits G_q_ via the RGS protein EAT-16 (Hajdu-Cronin *et al*. 1999). Thus, loss-of-function mutations in the channel pore-forming subunit NCA-1, its associated subunits UNC-79 and UNC-80, or the ER-localized NCA channel assembly factor NLF-1 suppress an activated G_q_ mutant, whereas a gain-of-function activating mutation in NCA-1 suppresses an activated G_o_ mutant. Though the NCA-1 channel has the greatest role in conveying G_q_-Rho signals, channels composed of the NCA-2 pore-forming subunit are also probably regulated by G_q_ because elimination of both NCA-1 and NCA-2 is required to fully suppress activated G_q_ and activated Rho mutants.

Mutations that eliminate the NCA channels, such as *unc-80* or the *nca-1 nca-2* double mutant, suppress the locomotion and body posture phenotypes of an activated Rho mutant (Figures 6-8), suggesting that NCA channels act downstream of Rho in a linear pathway. Additionally, channels composed of the pore-forming subunit NCA-1 may be the main target of G_q_-Rho signaling. First, loss-of-function mutations in *nca-1* alone, but not *nca-2*, partially suppress activated G_q_ and activated Rho mutants. Second, activating gain-of-function mutants of *nca-1* have phenotypes reminiscent of activated G_q_ and activated Rho mutants, but no activated mutants in *nca-2* have been isolated. Third, loss-of-function mutations in *nca-1*, but not *nca-2*, enhance the weaker fainting phenotype of an *nlf-1* mutant which does not fully eliminate function of the NCA channels. Together these data suggest that although either NCA-1 or NCA-2 activity is sufficient for wild-type locomotion, NCA-1 is likely to be the main target of G protein regulation.

We characterized three locomotory behaviors in this manuscript: locomotion rate, waveform, and fainting. The G_q_-Rho-NCA pathway regulates all three behaviors, but our genetic epistasis and cell-specific rescue experiments suggest that these behaviors are differentially regulated and involve at least partially distinct sets of neurons. First, the hyperactive locomotion and high-amplitude waveform phenotypes of the activated G_q_ mutant are genetically separable, since they are differentially suppressed by mutations in the PLCβ pathway and the Rho-NCA pathway. Mutations in the Rho-NCA pathway suppress both the locomotion rate and high-amplitude waveform of activated G_q_ whereas mutations in the PLCβ pathway suppress the locomotion rate but only very weakly suppress the high-amplitude waveform of activated G_q_. Thus, G_q_ acts through both the PLCβ pathway and the Rho-NCA pathway to regulate locomotion rate, but primarily through the Rho-NCA pathway to regulate the waveform of the animal. Second, we found that NLF-1 activity in head acetylcholine neurons is sufficient to fully rescue the fainting phenotype of an *nlf-1* mutant. However, inhibition of Rho in the head acetylcholine neurons did not suppress the high-amplitude waveform of the activated G_q_ mutant, but inhibition of Rho in all neurons did suppress. This suggests that Rho does not act solely in head acetylcholine neurons to regulate the waveform. Thus, the G_q_-Rho-NCA pathway acts in at least two different classes of neurons to regulate fainting and waveform.

It is not clear whether Rho activation of the NCA channels is direct or indirect. G_q_ directly interacts with and activates the RhoGEF and Rho, as shown by the crystal structure of a complex between G_q_α, RhoGEF, and Rho (Lutz *et al*. 2007). Rho is known to have many possible effectors and actions in cells (Etienne-Manneville and Hall 2002; Jaffe and Hall 2005). In *C. elegans* neurons, Rho has been previously shown to regulate synaptic transmission via at least two pathways, one involving a direct interaction of Rho with the DAG kinase DGK-1 and one that is DGK-1 independent (McMullan *et al*. 2006). A candidate for the link between Rho and NCA is the type I phosphatidylinositol 4-phosphate 5-kinase (PIP5K) that synthesizes the lipid phosphatidylinositol 4,5,-bisphosphate (PIP2). PIP5K is an intriguing candidate for several reasons. First, activation of G_q_ in mammalian cells has been shown to stimulate the membrane localization and activity of PIP5K via a mechanism that depends on Rho (Chatah and Abrams 2001; Weernink *et al*. 2004). Second, in *C. elegans*, mutations eliminating either the NCA channels (*nca-1 nca-2* double mutant) or their accessory subunits (*unc-79* or *unc-80*) suppress mutants in the PIP2 phosphatase synaptojanin (*unc-26*) and also suppress phenotypes caused by overexpression of the PIP5K gene *ppk-1* (Jospin *et al*. 2007). Loss of a PIP2 phosphatase or overexpression of a PIP5K are both predicted to increase levels of PIP2. Because the loss of NCA channels suppresses the effects of too much PIP2, it is possible that excessive PIP2 leads to overactivation of NCA channels and that PIP2 might be part of the normal activation mechanism. There are numerous examples of ion channels that are regulated or gated by phosphoinositide lipids such as PIP2 (Balla 2013), though PIP2 has not been shown to directly regulate NCA/NALCN.

The NCA/NALCN ion channel was discovered originally by bioinformatic sequence analyses (Lee *et al*. 1999; Littleton and Ganetzky 2000). It is conserved among all metazoan animals and is evolutionarily-related to the family of voltage-gated sodium and calcium channels (Liebeskind *et al*. 2012), forming a new branch in this super-family. Although the cellular role of the NCA/NALCN channel and how it is gated are not well understood, NALCN and its orthologs are expressed broadly in the nervous system in a number of organisms (Lee *et al*. 1999; Lear *et al*. 2005; Humphrey *et al*. 2007; Lu *et al*. 2007; Jospin *et al*. 2007; Yeh *et al*. 2008; Lu and Feng 2011; Lutas *et al*. 2016). Moreover, mutations in this channel or its auxiliary subunits lead to defects in rhythmic behaviors in multiple organisms (Lear *et al*. 2005, 2013; Lu *et al*. 2007; Jospin *et al*. 2007; Yeh *et al*. 2008; Pierce-Shimomura *et al*. 2008; Xie *et al*. 2013; Funato *et al*. 2016). Thus, NCA/NALCN is likely to play an important role in controlling membrane excitability. Additionally, NALCN currents have been reported to be activated by two different G protein-coupled receptors (GPCRs), the muscarinic acetylcholine receptor and the substance P receptor, albeit in a G protein-independent fashion (Lu *et al*. 2009; Swayne *et al*. 2009), and by low extracellular calcium via a G protein-dependent pathway (Lu *et al*. 2010). The latter study further showed that expression of an activated G_q_ mutant inhibited the NALCN sodium leak current, suggesting that high extracellular calcium tonically inhibits NALCN via a G_q_-dependent pathway and that low extracellular calcium activates NALCN by relieving this inhibition (Lu *et al*. 2010). By contrast, we find that G_q_ activates the NCA channels in *C. elegans*, and we show that the NCA channels are physiologically relevant targets of a G_q_ signaling pathway that acts through Rho.

In the last few years, both recessive and dominant human diseases characterized by a range of neurological symptoms including hypotonia, intellectual disability, and seizures have been shown to be caused by mutations in either *NALCN* or *UNC80* (Köroğlu *et al*. 2013; Al-Sayed *et al*. 2013; Chong *et al*. 2015; Aoyagi *et al*. 2015; Shamseldin *et al*. 2016; Stray-Pedersen *et al*. 2016; Gal *et al*. 2016; Fukai *et al*. 2016; Karakaya *et al*. 2016; Perez *et al*. 2016; Bend *et al*. 2016; Lozic *et al*. 2016; Valkanas *et al*. 2016; Wang *et al*. 2016; Vivero *et al*. 2017). Notably, dominant disease-causing mutations in the NALCN channel were modeled in worms and resemble either dominant activated or loss-of-function NCA mutants such as the ones we use in this study (Aoyagi *et al*. 2015; Bend *et al*. 2016). Human mutations in other components of the pathway we have described may cause similar clinical phenotypes.

## Acknowledgments

We thank Shohei Mitani for the *nlf-1(tm3631)* mutant; Steve Nurrish for worm strains and plasmids carrying activated Rho or C3 transferase; Ken Miller for a plasmid with the *unc-17*β promoter; Wayne Davis for FLP/FRT plasmids; Yuji Kohara for cDNA clones; the Sanger Center for cosmids; Brooke Jarvie, Jill Hoyt, and Michelle Giarmarco for the isolation of *unc-79* and *unc-80* mutations in the G_q_ suppressor screen; and Dana Miller for the use of her microscope and camera to take worm photographs. Some strains were provided by the CGC, which is funded by NIH Office of Research Infrastructure Programs (P40 OD010440). M.A. is an Ellison Medical Foundation New Scholar. E.M.J. is an Investigator of the Howard Hughes Medical Institute. This work was supported by NIH grants R00 MH082109 to M.A and R01 NS034307 to E.M.J.

---

**Figure S1.**
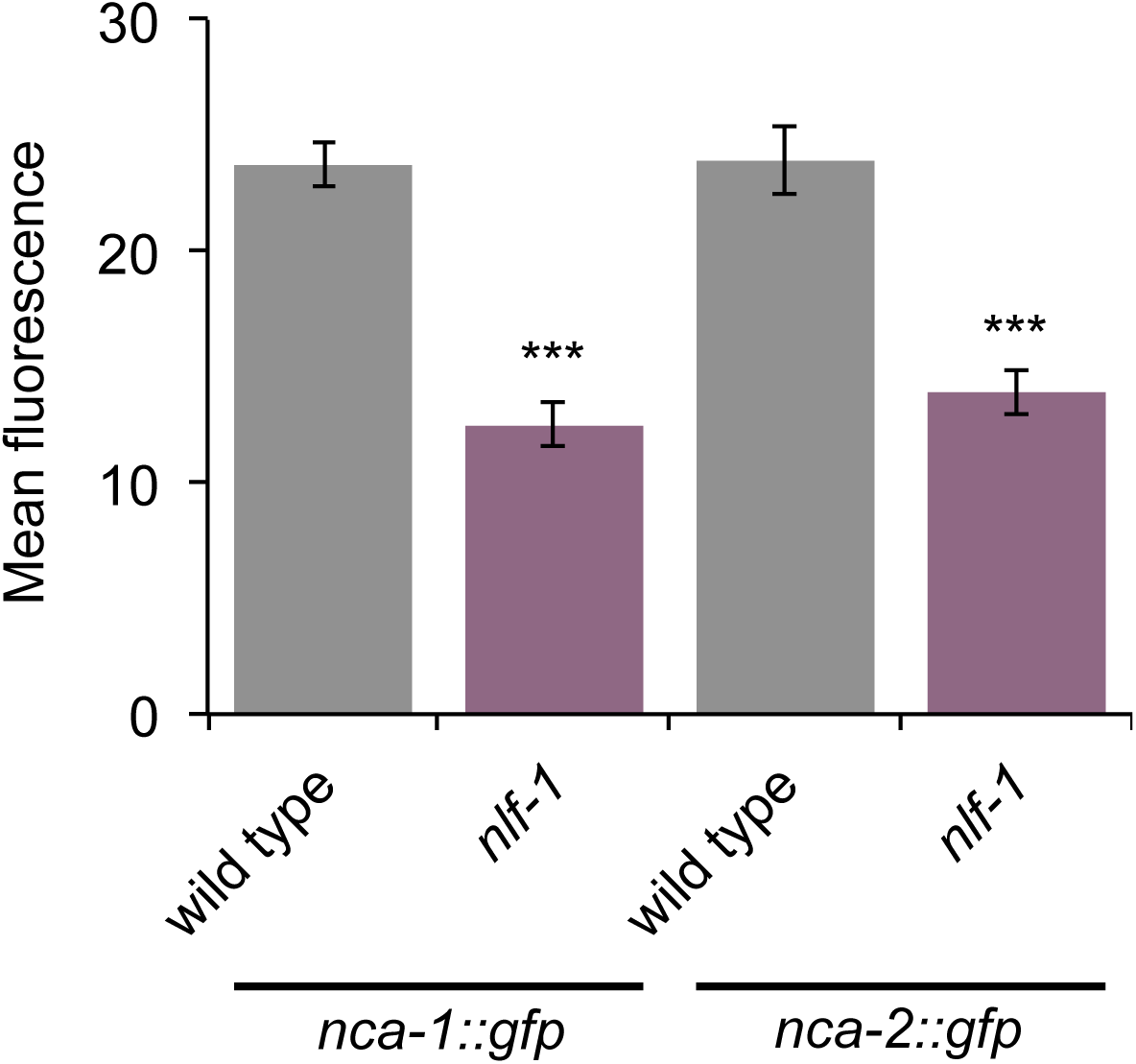
*nlf-1* mutants have reduced *nca-1::gfp* and *nca-2::gfp* fluorescence in the nerve ring. Quantification of the nerve ring fluorescence of C-terminally tagged *nca-1::gfp* (transgene *vaIs46*) and *nca-2::gfp* (transgene *vaIs41*) in wild type and *nlf-1(ox327)* mutants. ***, P<0.001, two-tailed unpaired t tests where each mutant strain is compared to its associated wild-type control. Error bars = SEM; n = 11-12.

**Table S1.**
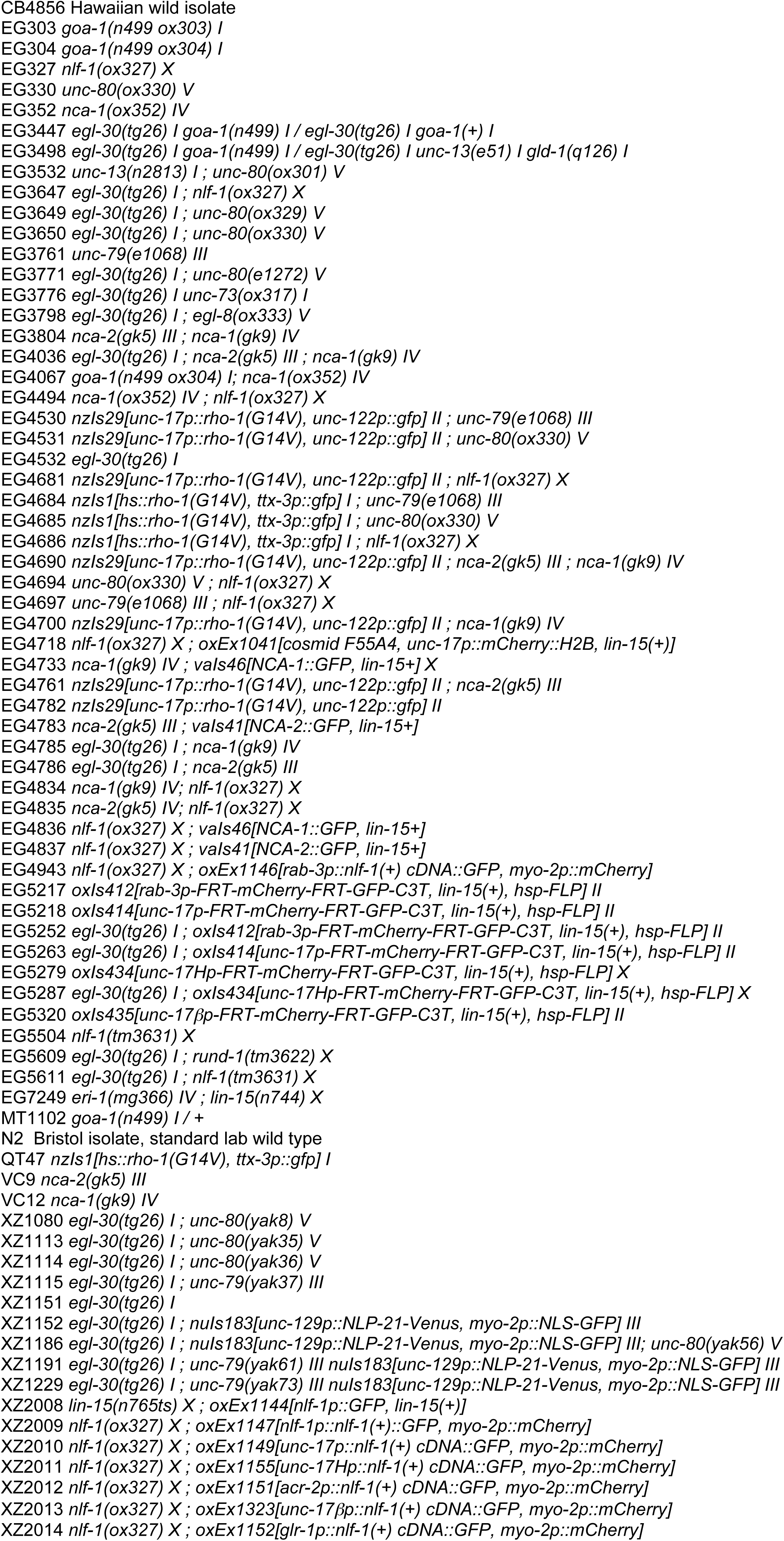
**List of strains**

**Table S2.**
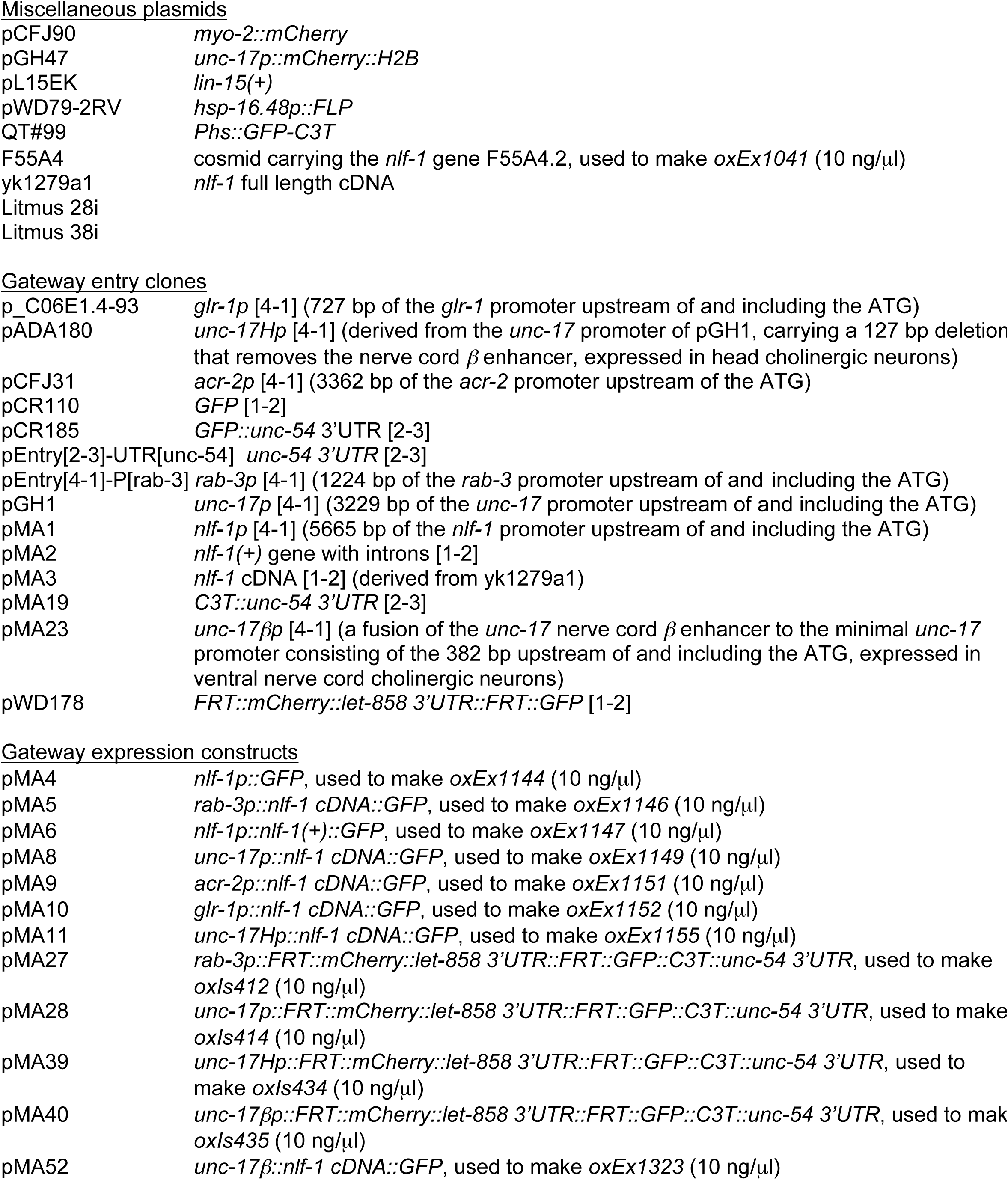
**List of plasmids**

|  |  |
| --- | --- |
| pCFJ90 | *myo-2::mCherry* |
| pGH47 | *unc-17p::mCherry::H2B* |
| pL15EK | *lin-15(+)* |
| pWD79-2RV | *hsp-16.48p::FLP* |
| QT#99 | *Phs::GFP-C3T* |
| F55A4 | cosmid carrying the *nlf-1* gene F55A4.2, used to make *oxEx1041* (10 ng/μl) |
| yk1279a1 | *nlf-1* full length cDNA |
| Litmus 28i |  |
| Litmus 38i |  |
| Gateway entry clones |  |
| p_C06E1.4-93 | *glr-1p* [4-1] (727 bp of the *glr-1* promoter upstream of and including the ATG) |
| pADA180 | *unc-17Hp* [4-1] (derived from the *unc-17* promoter of pGH1, carrying a 127 bp deletion that removes the nerve cord $\beta$ enhancer, expressed in head cholinergic neurons) |
| pCFJ31 | *acr-2p* [4-1] (3362 bp of the *acr-2* promoter upstream of the ATG) |
| pCR110 | *GFP* [1-2] |
| pCR185 | *GFP::unc-54* 3'UTR [2-3] |
| pEntry[2-3]-UTR[unc-54] | *unc-54* 3'UTR [2-3] |
| pEntry[4-1]-P[rab-3] | *rab-3p* [4-1] (1224 bp of the *rab-3* promoter upstream of and including the ATG) |
| pGH1 | *unc-17p* [4-1] (3229 bp of the *unc-17* promoter upstream of and including the ATG) |
| pMA1 | *nlf-1p* [4-1] (5665 bp of the *nlf-1* promoter upstream of and including the ATG) |
| pMA2 | *nlf-1(+)* gene with introns [1-2] |
| pMA3 | *nlf-1* cDNA [1-2] (derived from yk1279a1) |
| pMA19 | *C3T::unc-54* 3'UTR [2-3] |
| pMA23 | *unc-17βp* [4-1] (a fusion of the *unc-17* nerve cord $\beta$ enhancer to the minimal *unc-17* promoter consisting of the 382 bp upstream of and including the ATG, expressed in ventral nerve cord cholinergic neurons) |
| pWD178 | *FRT::mCherry::let-858* 3'UTR::*FRT::GFP* [1-2] |
| Gateway expression constructs |  |
| pMA4 | *nlf-1p::GFP*, used to make *oxEx1144* (10 ng/μl) |
| pMA5 | *rab-3p::nlf-1* cDNA::*GFP*, used to make *oxEx1146* (10 ng/μl) |
| pMA6 | *nlf-1p::nlf-1(+)*::*GFP*, used to make *oxEx1147* (10 ng/μl) |
| pMA8 | *unc-17p::nlf-1* cDNA::*GFP*, used to make *oxEx1149* (10 ng/μl) |
| pMA9 | *acr-2p::nlf-1* cDNA::*GFP*, used to make *oxEx1151* (10 ng/μl) |
| pMA10 | *glr-1p::nlf-1* cDNA::*GFP*, used to make *oxEx1152* (10 ng/μl) |
| pMA11 | *unc-17Hp::nlf-1* cDNA::*GFP*, used to make *oxEx1155* (10 ng/μl) |
| pMA27 | *rab-3p::FRT::mCherry::let-858* 3'UTR::*FRT::GFP::C3T::unc-54* 3'UTR, used to make *oxIs412* (10 ng/μl) |
| pMA28 | *unc-17p::FRT::mCherry::let-858* 3'UTR::*FRT::GFP::C3T::unc-54* 3'UTR, used to make *oxIs414* (10 ng/μl) |
| pMA39 | *unc-17Hp::FRT::mCherry::let-858* 3'UTR::*FRT::GFP::C3T::unc-54* 3'UTR, used to make *oxIs434* (10 ng/μl) |
| pMA40 | *unc-17βp::FRT::mCherry::let-858* 3'UTR::*FRT::GFP::C3T::unc-54* 3'UTR, used to make *oxIs435* (10 ng/μl) |
| pMA52 | *unc-17β::nlf-1* cDNA::*GFP*, used to make *oxEx1323* (10 ng/μl) |

